# Membrane fission during bacterial spore development requires DNA-pumping driven cellular inflation

**DOI:** 10.1101/2021.10.08.463650

**Authors:** Ane Landajuela, Martha Braun, Alejandro Martínez-Calvo, Christopher D. A. Rodrigues, Carolina Gomis Perez, Thierry Doan, David Z. Rudner, Ned S. Wingreen, Erdem Karatekin

**Author notes:** These authors contributed equally (Ane Landajuela, Martha Braun).

## Abstract

Bacteria require membrane fission for cell division and endospore formation. FisB catalyzes membrane fission during sporulation, but the molecular basis is unclear as it cannot remodel membranes by itself. Sporulation initiates with an asymmetric division that generates a large mother cell and a smaller forespore that contains only 1/4 of its complete genome. As the mother cell membranes engulf the forespore, a DNA translocase pumps the rest of the chromosome into the small forespore compartment, inflating it due to increased turgor. When the engulfing membranes undergo fission, the forespore is released into the mother cell cytoplasm. Here we show that forespore inflation and FisB accumulation are both required for efficient membrane fission. We suggest that high membrane tension in the engulfment membrane caused by forespore inflation drives FisB-catalyzed membrane fission. Collectively our data indicate that DNA-translocation has a previously unappreciated second function in energizing FisB-mediated membrane fission under energy-limited conditions.

**HIGHLIGHTS:**

- Membrane fission during endospore formation requires rapid forespore inflation by ATP-driven DNA translocation.
- Forespore inflation is fast enough to increase the tension of the engulfment and forespore membranes to near lysis tensions.
- FisB catalyzes membrane fission by impeding the flux of lipids that partially support forespore and engulfment membrane growth.
- Membrane fission utilizes chemical energy transduced to mechanical energy during DNA-packing into the forespore.

## INTRODUCTION

When nutrients are scarce, spore-forming bacteria initiate a morphological process called sporulation, which produces dormant, stress-resistant spores that can remain dormant for many years until conditions are favorable for germination (Moir and Cooper, 2015; Setlow, 2003). The first morphological event during sporulation is the formation of an asymmetric septum that produces a small forespore and a larger mother cell (Figure 1A). The mother cell membranes then engulf the forespore in a process reminiscent of phagocytosis. When the edges of the engulfing membrane reach the cell pole, they undergo membrane fission to release the forespore (Figure 1A inset), now surrounded by two membranes, into the mother-cell cytoplasm. The mother cell then packages the forespore in a protective cortex and coat while the forespore prepares for dormancy. When the forespore is mature, the mother cell lyses, releasing the dormant spore into the environment. Before asymmetric division, the chromosomes are remodeled into an axial filament with the origins of replication at the cell poles and the termini at the mid-cell (Webb et al., 1997). Thus, immediately after asymmetric division, only ~1/4 of the forespore chromosome is trapped inside the forespore (Wu and Errington, 1994, 1998). The rest of the chromosome is pumped into the forespore by an ATPase, SpoIIIE (Bath et al., 2000; Wu and Errington, 1997), consuming one ATP molecule per every two base pairs translocated (Liu et al., 2015). As the chromosome is packed into the forespore, the forespore volume doubles from its initial volume of ~0.1 μm^3^ by the time the engulfment membrane migrates completely around the forespore and continues to grow to ~0.3 μm^3^ about 3 h after septation (Lopez-Garrido et al., 2018). Forespore inflation is driven by osmotic forces which are mainly due to the counterions that are needed to neutralize the highly charged DNA (Lopez-Garrido et al., 2018). Upon complete DNA packing, the pressure difference between the forespore and mother cell reaches ~60 kPa (Lopez-Garrido et al., 2018). It has been suggested that DNA translocation can be seen as an energy transduction process, converting the chemical energy stored in ~1.5 million ATP molecules to mechanical energy, stretching the thin peptidoglycan layer between the forespore and engulfment membranes and smoothing their wrinkles (Lopez-Garrido et al., 2018). However, it is not clear if this stored mechanical energy is ever utilized by the nutrient-deprived, sporulating bacteria, and if so, how.

**Figure 1.**
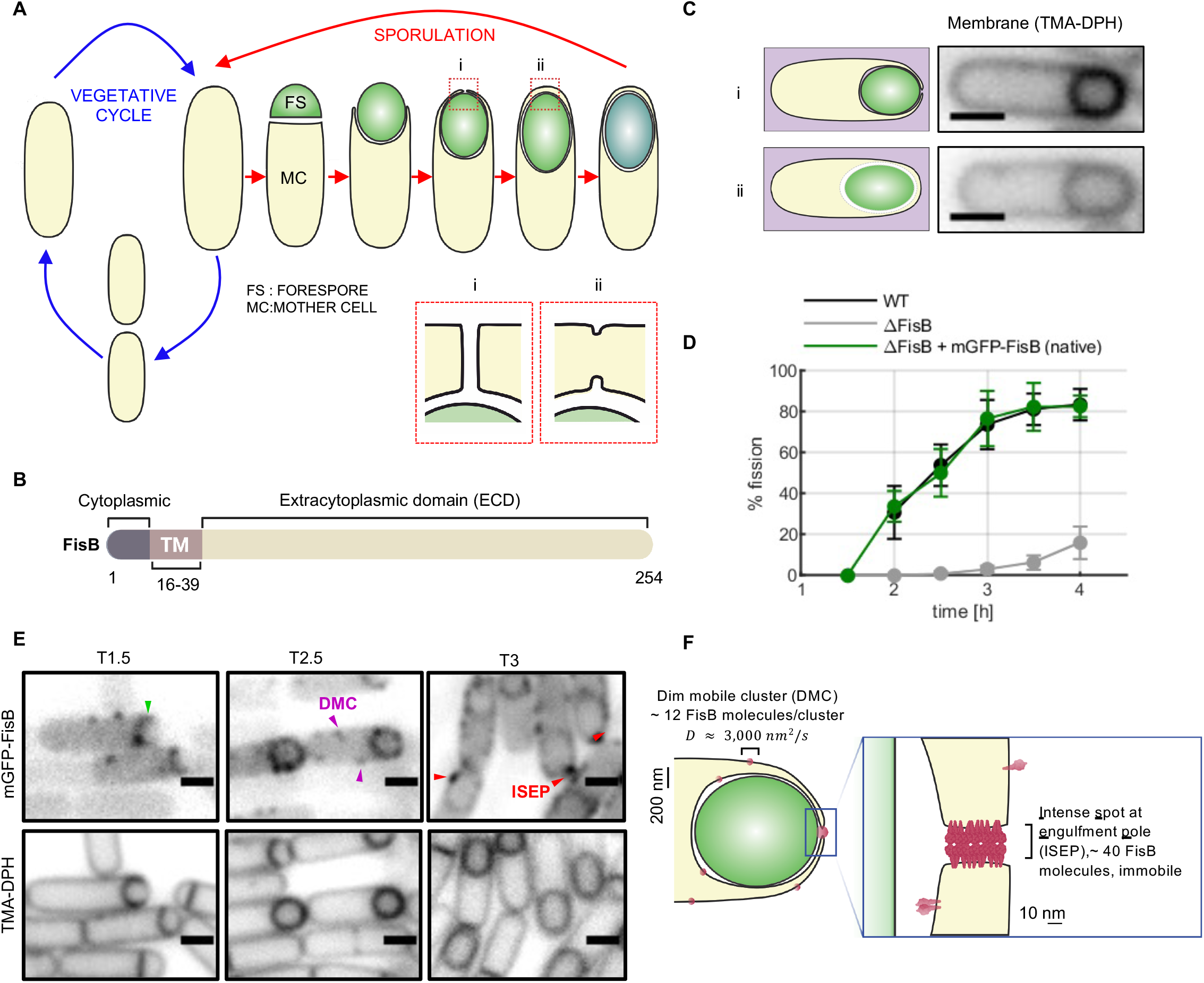
FisB is required for efficient membrane fission during sporulation. **A.** Schematic of sporulation stages. Upon starvation, sporulation is initiated by asymmetric division into a mother cell (MC) and a forespore (FS). The MC then engulf the FS. At the end of engulfment, a thin neck or tube connects the engulfment membrane to the rest of the MC membrane (red box). Fission of the neck releases the FS, now surrounded by two membranes, into the MC cytoplasm. Upon maturation, the FS turns into a spore and the MC lyses to release it. Inset: membrane fission step marking the end of engulfment. **B**. Domain structure of FisB (Doan et al., 2013; Landajuela et al., 2021). **C**. Membrane fission assay. An aliquot of cells is labeled with the lipophilic dye 1-(4-trimethylammoniumphenyl)-6-phenyl-1,3,5-hexatriene p-toluenesulfonate (TMA-DPH) which fluoresces only once inserted into the membrane. The dye labels internal membranes poorly. If a cell has not yet undergone membrane fission at the time of TMA-DPH labeling, the dye has access to the space between the engulfment and FS membranes, resulting in brighter labeling of the MC, FS, and engulfment membranes when these are adjacent to one another (top row). After fission, the dye labels internal membranes poorly (bottom row). **D**. Time course of membrane fission for wild-type cells, *ΔfisB* cells, or *ΔfisB* cells complemented with mGFP-FisB expressed at native levels (strain BAM003). Some membrane fission occurs even in the absence of FisB (*ΔfisB*, gray markers). Mean±SEM from 3 independent experiments are shown (>300 cells were analyzed per point). **E.** Wide-field fluorescence microscopy images of cells expressing mGFP-FisB at native levels (strain BAM003). Aliquots were taken from the suspension at indicated times after nutrient downshift, labeled with TMA-DPH, and images of mGFP-FisB and membranes were acquired sequentially. Examples of sporulating cells with mGFP-FisB enriched at the septum (1.5 h), forming a dim mobile cluster (DMC; 2.5 h) and with a discrete mGFP-FisB focus at the cell pole (intense spot at engulfment pole, ISEP, 3 h) are highlighted with arrowheads. Scale bars represent 1 μm. **F**. Schematic of FisB dynamics (Landajuela et al., 2021). At the end of engulfment, ~40 FisB molecules accumulate at the neck into an immobile cluster to catalyze membrane fission.

Here, we investigate the relationship between forespore inflation driven by DNA translocation and the membrane fission event that releases the forespore into the mother cell cytoplasm. We previously showed that this event is catalyzed by the protein FisB that accumulates at the neck of the engulfing membranes during sporulation in *Bacillus subtilis* (Doan et al., 2013; Landajuela et al., 2021). FisB, a ~250 residue protein that is conserved among endospore-forming bacteria, is produced after asymmetric division in the mother cell (Doan et al., 2013). It possesses a short N-terminal cytoplasmic domain, a single-pass transmembrane domain (TMD), and a large extracytoplasmic domain (ECD, Figure 1B) (Landajuela et al., 2021). In cells lacking FisB (*ΔfisB*), engulfment proceeds normally, but the membrane fission step is impaired (Doan et al., 2013; Landajuela et al., 2021). Membrane fission can be assessed during a sporulation time course by labeling cells with a lipophilic dye that crosses the cell membrane inefficiently (Figure 1C). In cells that have not yet undergone fission, the dye has access to the forespore, engulfing, and mother cell membranes, resulting in a stronger signal where these three membranes are in close proximity. By contrast, in cells that have undergone membrane fission, the dye only weakly labels the internal membranes, allowing the quantification of cells that have undergone fission at a given time point after initiation of sporulation (Landajuela et al., 2021) (Figure 1D). A functional GFP-FisB fusion forms dim mobile clusters (DMC), each containing ~12 GFP-FisB molecules. An intense spot at the engulfment pole (ISEP) containing ~40 copies of GFP-FisB appears around the time of membrane fission (Figure 1E,F and (Landajuela et al., 2021)). FisB localizes to the membrane neck that connects the engulfment membrane to the rest of the mother cell membrane based only on lipid-binding, self-aggregation, and the unique geometry encountered at the end of engulfment (Landajuela et al., 2021). FisB self-oligomerizes and binds acidic lipids, bridging artificial membranes, but does not appear to have other binding partners. During sporulation, the only location where FisB can bridge membranes is at the membrane neck that eventually undergoes membrane fission (Figure 1F), suggesting homo-oligomerization and trans interactions that bridge membranes are sufficient to explain FisB accumulation in the neck during late stages of engulfment (Landajuela et al., 2021). However, how membrane fission occurs at the last step of engulfment is not known.

Here we show that forespore inflation and FisB accumulation at the membrane neck are both required for efficient membrane fission. Interventions that slowed or prevented forespore area expansion delayed or inhibited membrane fission, even when FisB had accumulated at the membrane neck. Conversely, sporulating cells lacking FisB had impaired membrane fission but their forespores expanded. We find that membrane fission is driven by the combination of high tension in the engulfment membrane due to forespore inflation and FisB oligomerization impeding lipid flux that partially supports forespore and engulfment membrane growth. Our results demonstrate that part of the mechanical energy stored during the process of DNA packaging into the forespore is used for membrane fission.

## RESULTS

### FisB does not remodel membranes, but forms a stable, extended network on GUV membranes

The simplest mechanism of FisB-catalyzed membrane fission would involve direct membrane remodeling by FisB, similar to eukaryotic membrane fission machineries dynamin (Ferguson and De Camilli, 2012) or ESCRT-III (Schoneberg et al., 2017). To test this idea, we studied interactions of the FisB extracellular domain (ECD) with model GUV membranes. We fluorescently labeled recombinant purified FisB ECD with iFluor555-maleimide at a cysteine introduced at residue 123 (iFluor-FisB ECD, Figure 2A), a substitution that does not affect FisB’s function (Landajuela et al., 2021). GUV membranes were labeled by including a fluorescent lipid in the lipid composition. Membranes and the labeled protein were then visualized using spinning-disc confocal (SDC) microscopy (Figure 2). We first incubated GUVs with various concentrations of iFluor-FisB ECD in a closed chamber for 60 min, then imaged the membranes and FisB ECD (Figure 2B). iFluor-FisB ECD did not bind GUVs lacking acidic lipids, as reported previously (Landajuela et al., 2021). By contrast, iFluor-FisB ECD bound GUVs containing 30 mole % cardiolipin (CL). At 20 nM in solution, FisB binding to GUVs was barely detectable, but at ~75 nM, discrete FisB spots were visible. These spots diffused around the GUV membrane, reminiscent of the DMC in live cells. At 200 nM and above, most GUVs were covered more uniformly and completely by iFluor-FisB ECD. Consistent with previous work (Landajuela et al., 2021), and unlike many proteins implicated in eukaryotic membrane fission (Campelo and Malhotra, 2012; Renard et al., 2018; Simunovic et al., 2019), FisB did not cause any membrane deformations such as tubulation or invaginations. Because such deformations are opposed by membrane tension, we repeated these experiments using deflated GUVs with lower membrane tension. Even at high coverage and when the GUVs were deflated, FisB did not cause any remodeling of GUV membranes (Figure 2B). However, we noticed that deflated GUVs covered with iFluor-FisB ECD did not display shape fluctuations typical of bare deflated GUVs, suggesting FisB may form a fixed, extended network on the membranes stabilizing the membrane shape.

**Figure 2.**
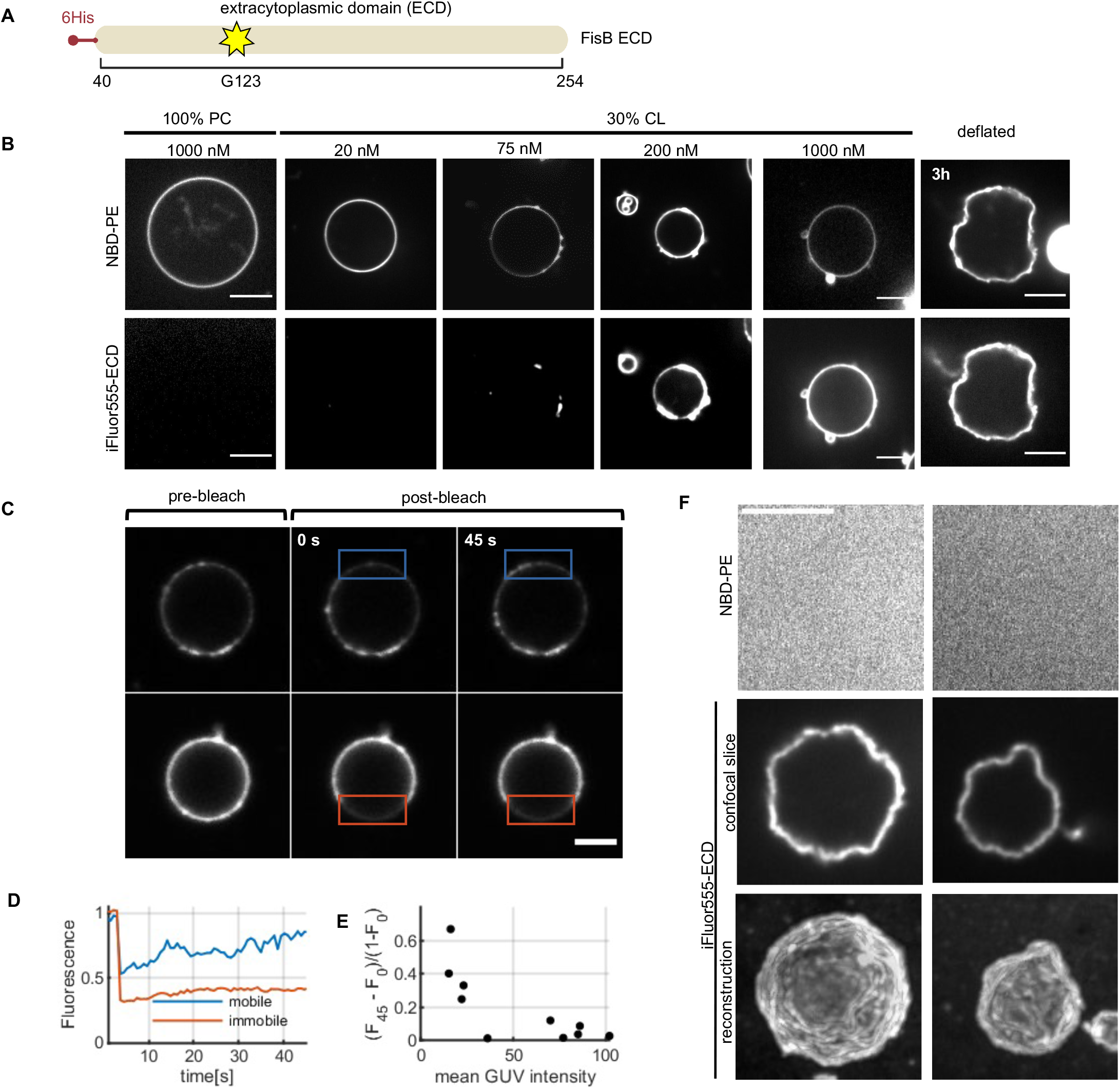
FisB does not remodel membranes. **A**. The soluble recombinant fragment of FisB comprising its extracytoplasmic domain (ECD) used in the experiments with GUVs. **B**. Interactions of FisB ECD with GUV membranes (“100% PC” and “30% CL” membranes are composed of (all mole %) 99 PC, 1 NBD-PE, and 30 *E. coli* CL, 69 eggPC, 1 NBD-PE, respectively). At low concentrations, FisB ECD forms small mobile clusters. With increasing concentration, labeling becomes more uniform. Even at high coverage or with deflated GUVs (right), no membrane remodeling is evident (deflated GUVs were composed of (all mole %) 25 *E. coli* PE, 5 *E. coli* CL, 50 *E. coli* PG, 19 eggPC and 1 DiD or NBD-PE (BS mix)). **C**. Mobility of FisB ECD decreases with increasing coverage. GUVs composed of BS mix were incubated with 1 μM iFluor555-FisB ECD for 2h. GUVs were covered with protein to varying degrees. Top row shows fluorescence recovery of a GUV with low iFluor555-FisB ECD coverage. Bottom row shows an example of a GUV fully covered with iFluor555-FisB ECD fluorescence that does not recover after photobleaching. Photobleached regions are indicated with boxes. **D**. Normalized fluorescence intensity from the boxed regions in C as a function of time. **E**. Higher protein coverage (*∝* mean GUV membrane intensity) leads to lower protein mobility. The fractional fluorescence recovery 45 s post-bleaching ((*F*_45_ – *F*_0_)/(1 – *F*_0_), where *F*_45_ is the fluorescence intensity at 45 s and *F*_0_ is the intensity just after bleaching, both relative to pre-bleach intensity) is plotted as a function of mean GUV membrane intensity. Each dot represents a GUV. **F**. FisB ECD forms a fixed, stable network on the GUV membrane that persists even after the membranes are dissolved with detergent. 1 μM iFluor555-FisB ECD was incubated with deflated GUVs for 2h, then Triton X-100 was added (1.7 mM final concentration, 7-8-fold above the critical micelle concentration (Tiller et al., 1984)), and the sample imaged 5-10 min thereafter.

**Figure 3.**
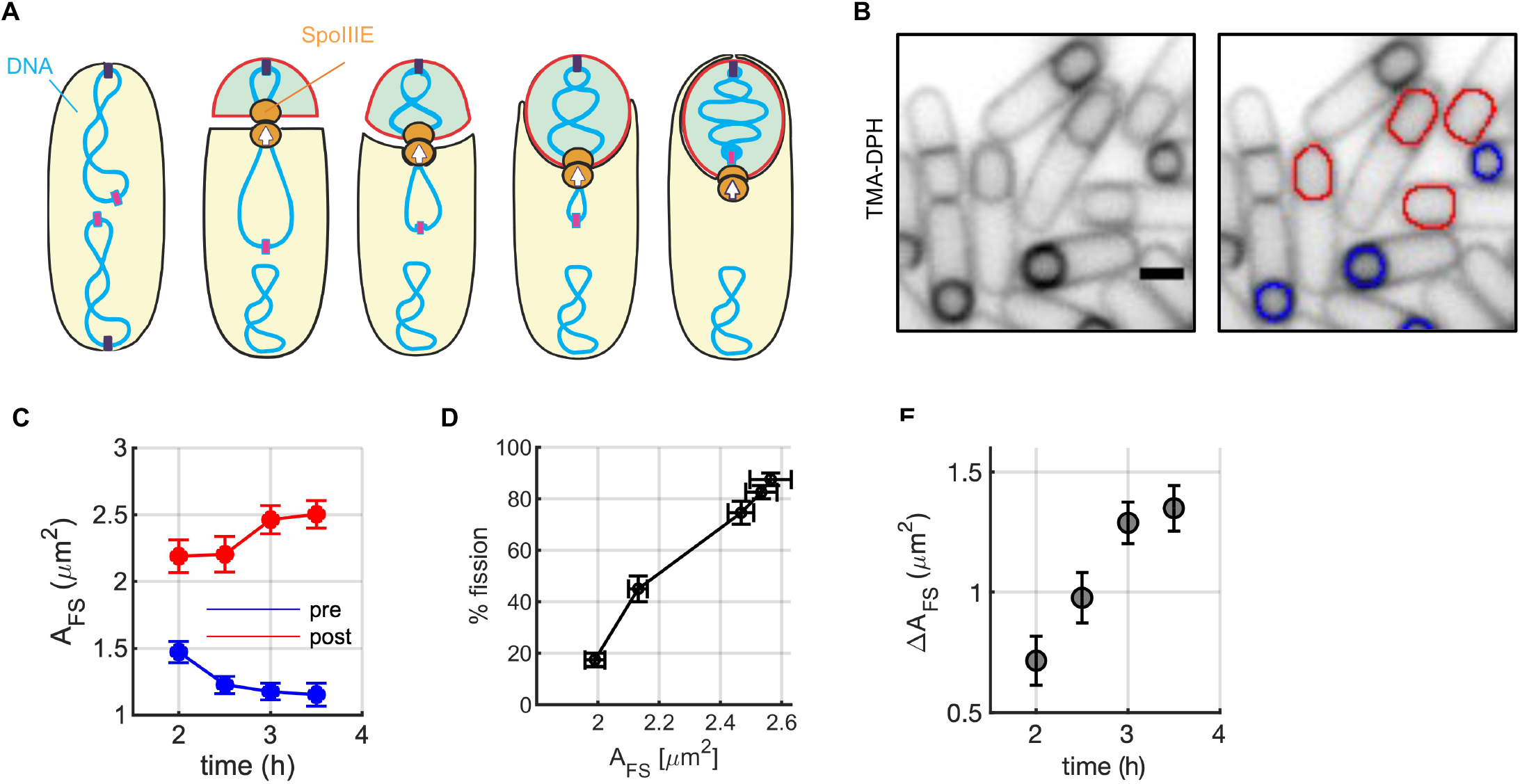
Forespore area increase during sporulation correlates with membrane fission activity. **A**. Schematic of SpoIIIE-mediated pumping of the chromosome into the forespore leading to forespore inflation. **B**. Images show wild-type strain (PY79) at *t*=3 h after nutrient downshift to initiate sporulation. Membranes were visualized using TMA-DPH. Forespores of cells that have undergone membrane fission (red contours on the right panel) are larger than those that have not yet undergone membrane fission (blue contours). Scale bars represent 1 μm. **C**. Quantification of forespore membrane area for cells that have (red, “post”) or have not (blue, “pre”) undergone membrane fission. **D**. The percentage of cells that have undergone membrane fission as a function of post-fission forespore area. **E**. The difference between post- and pre-fission forespore areas, *ΔA*_FS_, grows as a function of time. In C-E, mean±SEM of 3 independent experiments are shown, with 70 cells analyzed per data point.

We reasoned that if FisB ECD does indeed form an extended network on membranes, its mobility should be low in such regions. To test this, we performed fluorescence recovery after photobleaching (FRAP) experiments (Figure 2C). We incubated 1 μM iFluor-FisB ECD with GUVs for 2 h. This allowed for a range of iFluor-FisB ECD coverage of GUVs. Some GUVs were uniformly covered by iFluor-FisB ECD whereas some others were covered only partially by iFluor-FisB ECD patches. We bleached the iFluor-FisB ECD fluorescence in a rectangular region of interest (ROI) and monitored the subsequent recovery on a given GUV. Recovery was faster for GUVs that had partial, patchy iFluor-FisB ECD coverage, as new patches diffused into the bleached region (Figure 2C upper row and Figure 2D). By contrast, there was virtually no recovery for GUVs that were uniformly covered by iFluor-FisB ECD (Figure 2C lower row and Figure 2D). To relate FisB ECD membrane coverage to mobility, we plotted the mean iFluor-FisB ECD signal along the contour of the GUV, 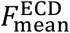, against the fractional fluorescence recovery 45 s after bleaching, as shown in Figure 2E. This analysis showed that increasing FisB ECD coverage led to decreased FisB ECD mobility, consistent with the idea that FisB ECD forms a denser, less dynamic network as coverage increases.

To test how stable the FisB ECD network is, we dissolved the membranes after the network had formed while observing iFluor-FisB ECD signals. Five to ten minutes after addition of Triton X-100 the lipid signal disappeared, suggesting that the membranes were dissolved. Remarkably, iFluor-FisB ECD signals were intact (Figure 2F), indicating that once formed, the three-dimensional FisB ECD network was stable even after the removal of the membrane.

Altogether, these data argue that FisB does not remodel membranes but is capable of forming an extended, stable network in which individual FisB molecules are immobile.

### Forespore inflation accompanies membrane fission

How can FisB catalyze membrane fission if it cannot remodel membranes by itself? We reasoned that an additional cellular process should be involved, given that direct interactions with other proteins could not be detected (Doan et al., 2013; Landajuela et al., 2021). A process that could potentially influence membrane fission is the forespore volume increase that occurs after asymmetric division due to translocation of DNA into the forespore by the ATPase SpoIIIE (Lopez-Garrido et al., 2018) (Figure 3A). However, whether and how this volume increase is related to membrane fission was not known. We began by investigating whether there was a correlation between forespore inflation and membrane fission. We used the lipophilic fluorescent dye 1-(4-trimethylammoniumphenyl)-6-phenyl-1,3,5-hexatriene p-toluenesulfonate (TMA-DPH) that allowed us to detect forespore contours of both cells that had undergone membrane fission and those that had not (Landajuela et al., 2021). We used a semi-automated active-contour fitting algorithm (Smith et al., 2010) and estimated the membrane areas from the surface of revolution around the long axis of symmetry (Methods). Three hours after initiation of sporulation via nutrient downshift, post-fission forespores were visually larger than their remaining pre-fission counterparts (Figure 3B). The average areas of pre- and post-fission forespores decreased and increased as a function of time after the nutrient downshift, respectively (Figure 3C), suggesting cells with rapidly growing forespores underwent membrane fission, leaving behind smaller pre-fission forespores. Similar trends were obtained using a soluble CFP forespore marker (Figure S1), ruling out potential artifacts due to the use of a lipophilic membrane dye. The close link between forespore inflation and membrane fission was even more evident when we plotted the percentage of cells having undergone membrane fission against the post-fission forespore area, which yielded a nearly linear relationship (Figure 3D).

The forespore area difference between pre- and post-fission cells, *ΔA*_FS_, increased during sporulation to reach 1.3 ± 0.1 μm^2^ (Figure 3E). Taking into account the 4 membrane leaflets that must grow together and assuming an area per lipid of ~0.7 nm^2^, ~7.4 × 10^6^ new lipids must be added to the engulfment and forespore membranes during sporulation. Given that this growth occurs over ~2 h, lipids must be added at an average rate of ~1,000 lipids/s to the forespore and engulfment membranes. Interestingly, vegetative cell membrane areas grow at a comparable rate (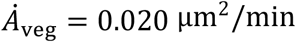, or about ~1000 lipids/s, taking into account two leaflets, 95% confidence interval: 0.007-0.023 μm^2^/min, n=8 cells), consistent with the finding that phospholipid synthesis during sporulation occurs near the maximal rate associated with vegetative growth (Pedrido et al., 2013). Thus, forespore inflation requires high rates of *de novo* phospholipid synthesis despite energy-limiting conditions, and most of the newly synthesized lipids are utilized for forespore area growth during the late stages of engulfment and after fission.

In summary, there is a clear correlation between membrane fission at the end of engulfment and forespore inflation, suggesting that forespore growth may facilitate membrane fission.

### Hypothesis: increasing membrane tension during forespore inflation drives FisB-dependent membrane fission

How can forespore inflation promote FisB-mediated membrane fission? We hypothesized that increased membrane tension due to forespore inflation, coupled with FisB oligomerization at the membrane neck, could work synergistically to drive membrane fission. Because DNA pumping into the forespore leads to forespore inflation through increased turgor, and smooths membrane wrinkles (Lopez-Garrido et al., 2018), there must be an associated increase in the tension of the forespore membrane. Furthermore, because the forespore and engulfment membranes are separated through a thin layer of peptidoglycan only ~20 nm wide (Khanna et al., 2019; Tocheva et al., 2011), we expect the tension and the area of the engulfment membrane (which eventually undergoes fission) to increase together with that of the forespore membrane. The membrane tension of the forespore and engulfment membranes can be estimated from the excess osmotic pressure in the forespore, ~60 kPa (Lopez-Garrido et al., 2018). Assuming the two membranes that surround the forespore balance the excess osmotic pressure, then according to Laplace’s Law Δ*P* = 4*γ*/*R*_fs_, so that a forespore radius of *R*_fs_ = 400 nm implies a membrane tension *γ* ~ 6 pN nm^-1^. This is a very high value, around values expected for membrane rupture (Evans et al., 2003; Needham and Nunn, 1990; Ojkic et al., 2014). It is thus likely that the peptidoglycan cell wall surrounding the forespore may actually be necessary during forespore inflation to avoid membrane rupture (Lopez-Garrido et al., 2018).

The hypothesis that increasing membrane tension during forespore inflation drives FisB-dependent membrane fission leads to the following predictions. First, interfering with forespore inflation should hinder membrane fission. Even when forespore inflation is slowed or blocked, FisB should be able to properly localize to the membrane neck (provided engulfment is not compromised), because FisB’s localization relies solely on its interactions with itself and with membranes, and the geometry of the neck (Landajuela et al., 2021). Second, in the absence of FisB, forespores should inflate but membrane fission should be impaired.

### Blocking lipid synthesis impedes forespore inflation and membrane fission

Lipid synthesis is required during vegetative growth (Yao et al., 2012; Zhang and Rock, 2008) and sporulation (Pedrido et al., 2013; Schujman et al., 1998) and fatty acid availability determines average cell size (Vadia et al., 2017). When *B. subtilis* cells are placed in a nutrient poor medium, lipid synthesis is initially downregulated, but with the onset of sporulation, it returns close to the maximal rate (Pedrido et al., 2013). We reasoned that blocking lipid synthesis at different time points after the initiation of sporulation should block forespore area expansion at different stages. Furthermore, if forespore inflation is needed for membrane fission, the percentage of forespores blocked in inflation should correlate with those stalled at the pre-fission stage.

We blocked the synthesis of neutral lipids and phospholipids at different time points *t* after nutrient downshift using cerulenin, a drug that inhibits *de novo* fatty acid biosynthesis (Heath and Rock, 2004) (Figure S2A). We then probed the cells for membrane fission at *t* = 3 h using TMA-DPH labeling. We found that the earlier lipid synthesis was blocked, the smaller the fraction of cells that had undergone membrane fission at *t* = 3 h (Figure S2B,C). Both pre-and post-fission forespore membrane areas decreased with longer cerulenin application (Figure S2D). These results are consistent with previous reports that *de novo* lipid synthesis is needed for engulfment and forespore inflation (Lopez-Garrido et al., 2018; Pedrido et al., 2013). In addition to perturbing engulfment (Figure S2E,F) FisB accumulation at the fission site was reduced in the presence of cerulenin (Figure S2G), making it difficult to infer the role of forespore inflation in membrane fission. In order to unmask the contribution of forespore inflation on membrane fission, we focused on cells in which FisB had successfully accumulated at the membrane fission site. We discovered that an increasing fraction of these cells failed to undergo membrane fission with longer cerulenin application (Figure S2H). Thus, although impaired engulfment and FisB accumulation at the fission site partially account for the membrane fission defects observed, blocking lipid synthesis inhibited membrane fission even for cells with proper FisB localization, suggesting forespore inflation facilitates membrane fission.

### Slowing DNA translocation slows membrane fission

In addition to lipid synthesis, forespore inflation requires ATP-dependent DNA translocation into the forespore. Accordingly, in a complementary set of experiments we took advantage of previously characterized mutations in the DNA translocase SpoIIIE to investigate whether forespore inflation contributes to membrane fission.

We began by using an ATPase mutant (SpoIIIE36) that is translocation defective (Pogliano et al., 1997). In the absence of DNA pumping, the forespores remained small and engulfment was severely perturbed (Figure S3A), similar to the engulfment phenotype of the *ΔspoIIIE* mutant reported previously (Doan et al., 2013). Membrane fission was severely impaired in these cells (Figure S3B) as was forespore area growth, both for pre-fission and the small number of post-fission forespores (Figure S3C). The fission defect can only be partly explained by deficient FisB localization, as 23±4 % of the SpoIIIE36 cells had an ISEP at 3 h after the nutrient downshift, compared to 70±6% of WT cells (Figure S3G), yet only 5% of the SpoIIIE36 cells had undergone membrane fission at this timepoint (Figure S3B). Importantly, among cells with an ISEP at the end of engulfment, only ~16±4% of SpoIIIE36 cells had undergone membrane fission compared to ~91±6% % of WT cells (Figure S3H). This difference was even more pronounced when we used a sensitized background in which FisB was expressed at ~8-fold lower levels than wild-type (Doan et al., 2013; Landajuela et al., 2021). Under these conditions, there were no post-fission SpoIIIE36 cells at hour 3 of sporulation among the 7% that had an ISEP. By contrast, 95% of the sporulating cells with SpoIIIE^WT^ that had an ISEP had undergone membrane fission in this FisB low-expression background (Figure S3G,H). Altogether, these results suggest that membrane fission requires both FisB accumulation at the neck (ISEP formation) and forespore inflation.

The severe engulfment defects in the SpoIIIE36 mutant have the potential to obscure the contribution of forespore inflation to FisB localization and membrane fission. Accordingly, we took advantage of a second mutant (SpoIIIE^D584A^) that translocates DNA ~2.5-fold slower than wild-type (Burton et al., 2007) and does not have engulfment defects (Figure 4). In sporulating cells expressing *spoIIIE*^D584A^, forespore inflation was delayed for post-fission cells and the average forespore area of the remaining pre-fission cells increased instead of decreasing as for wild-type cells, suggesting slow forespore inflation slows membrane fission (Figure 4A-C). This was more evident when the percentage of cells that had undergone fission was plotted together with the post-fission forespore areas as functions of time (Figure 4D). The percentage of cells that underwent membrane fission increased with the increase in post-fission forespore area (Figure 4E). Importantly, engulfment (Figure 4F,G) and FisB localization (Figure 4H) were similar to wild-type. In fact, by hour 3 of sporulation, *spoIIIE*^D584A^ cells accumulated ~1.4-fold more FisB at the cell pole than did wild-type cells (Figure 4I), yet only ~40% of the *spoIIIE*^D584A^ cells had undergone membrane fission compared to ~75% of the wild type cells (Figure 4D), demonstrating that accumulating more FisB does not necessarily lead to more efficient membrane fission.

**Figure 4.**
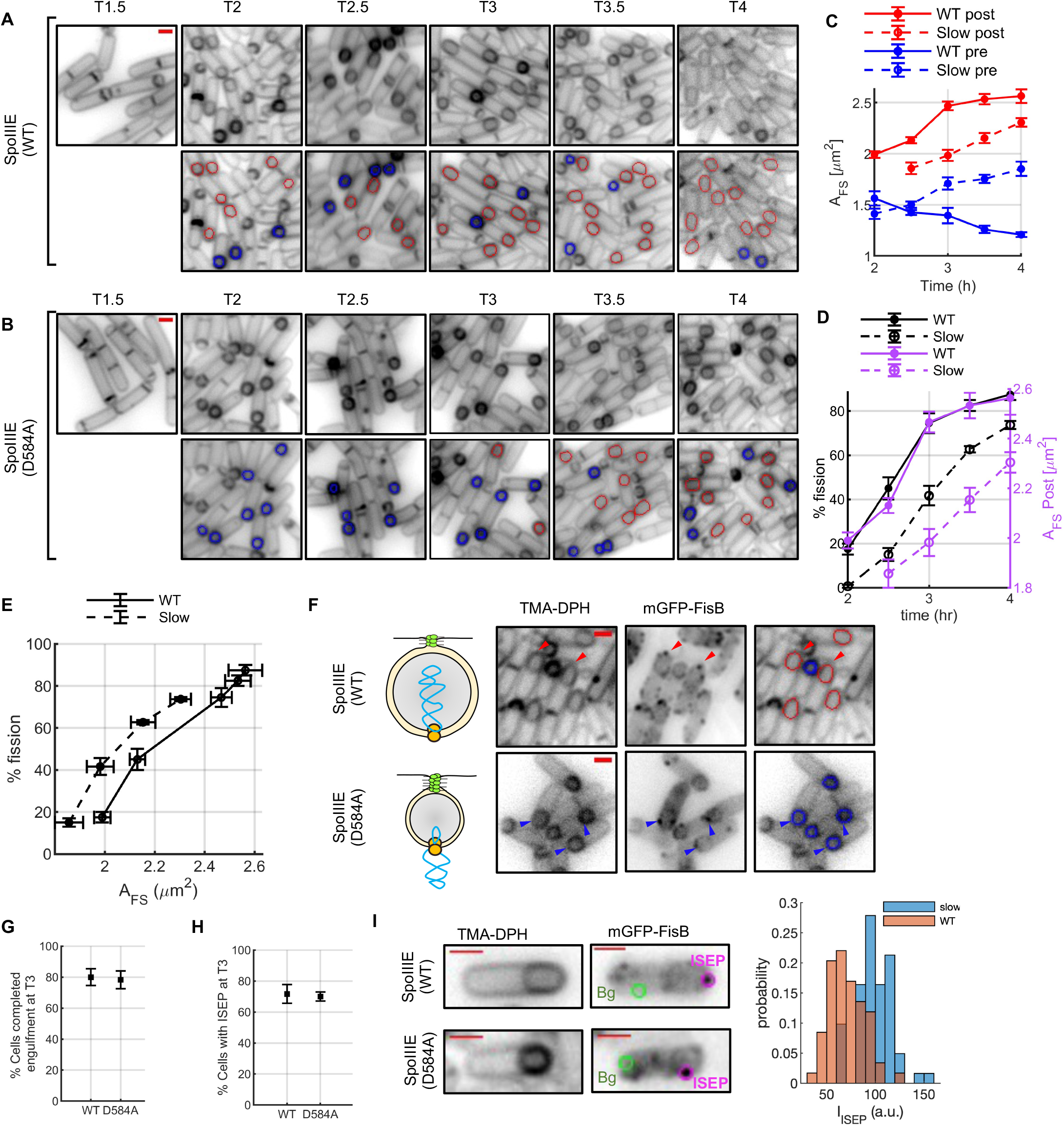
Slower forespore inflation leads to slower membrane fission with no adverse effect on engulfment or FisB localization. **A**. Fluorescence microscopy images of TMA-DPH labeled wild-type SpoIIIE strain BAL039 (top row, times after nutrient downshift are indicated). The lower row shows the same images, but with the overlaid forespore contours for cells that have (red) or have not (blue) undergone membrane fission. **B.** As in A, but for the slow DNA-pumping SpoIIIE strain BAL040 (SpoIIE^D584A^). **C**. The average forespore area for pre- (blue) and post-fission (red) cells as a function of time into sporulation for the SpoIIE^D584A^ (dashed lines, strain BAL040) or wild-type (solid lines) cells (strain BAL039). (n=25-70 cells per data point). **D**. The percentage of cells that have undergone membrane fission (left axis, black) followed a very similar time course as the forespore area as a function of time (right axis, magenta) for both WT (solid lines) and SpoIIE^D584A^ (dashed lines) cells. Both membrane fission and forespore inflation were delayed in SpoIIE^D584A^ cells (>300 cells per data point). **E**. The percentage of post-fission cells increased with increasing forespore area for both WT (black) and SpoIIE^D584A^ (magenta) cells. WT data is copied from Figure 3 for comparison. **F.** Fluorescence microscopy images of WT or SpoIIE^D584A^ cells expressing mGFP-FisB at native levels in a *ΔfisB* background, probed 3h after the nutrient downshift. Membranes were labeled with TMA-DPH. In cells expressing WT SpoIIIE (top row) post-fission cells have larger forespores (red contours) and FisB is accumulated at the membrane fission site (red arrowheads). At this stage, most cells expressing SpoIIE^D584A^ are still in the pre-fission stage, even if engulfment is visually complete and FisB has accumulated at the membrane fission site. **G.** Percentage of WT or SpoIIE^D584A^ cells with engulfment membrane migration completed by t=3 h into sporulation. Engulfment is not perturbed significantly by the slow DNA pumping by SpoIIE^D584A^ (>300 cells per data point). **H**. FisB localization is not perturbed in SpoIIE^D584A^ cells compared to WT cells. **I**. Distributions of mGFP-FisB fluorescence intensities for the ISEP in willd-type (“WT”) and the slow DNA pumping by SpoIIE^D584A^ mutant strain (“slow”). Representative regions of interest (and regions taken for background correction, “Bg”) are shown on the example images on the left. The means (± SD) are 70 ± 17 a.u. for WT and 99 ± 18 for SpoIIE^D584A^ ISEP intensities. Since for WT this intensity corresponds to ~40 copies of FisB (Landajuela et al., 2021), we conclude that nearly 60 copies are accumulated in SpoIIE^D584A^. Scale bars represent 1 μm in A, B, and F. Means±SEM for 3 biological replicates are plotted in panels C-E, G,H. See Figure S3 for results with an ATPase dead SpoIIIE mutant.

Together, these results strongly suggest that FisB accumulation at the membrane neck is *not* a sufficient condition for membrane fission; fission additionally requires forespore enlargement.

### In the absence of FisB, forespores inflate but fission is impaired

Our data suggest that FisB accumulation at the fission site and forespore inflation work together to drive membrane fission. To investigate whether these two processes occur independently, we monitored forespore membrane area changes in cells expressing FisB at low levels or altogether lacking it.

We analyzed forespore area at different time points after a nutrient downshift, using TMA-DPH (Figure 5). As anticipated, most sporulating cells lacking FisB (*ΔfisB*) did not undergo membrane fission (Figure 1D) despite completing engulfment (Figure 5A) (Doan et al., 2013; Landajuela et al., 2021). Remarkably, the forespore areas of these pre-fission cells, 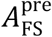, increased as a function of time after the nutrient downshift, nearly doubling by *t* = 3.5 h, while those of wild-type cells decreased slightly during the same period (Figure 5B). The forespore areas of the small fraction of *ΔfisB* cells that underwent membrane fission (Figure 1), 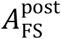, were slightly larger than those of wild-type cells and followed a similar time course, increasing as a function of time (Figure 5C).

**Figure 5.**
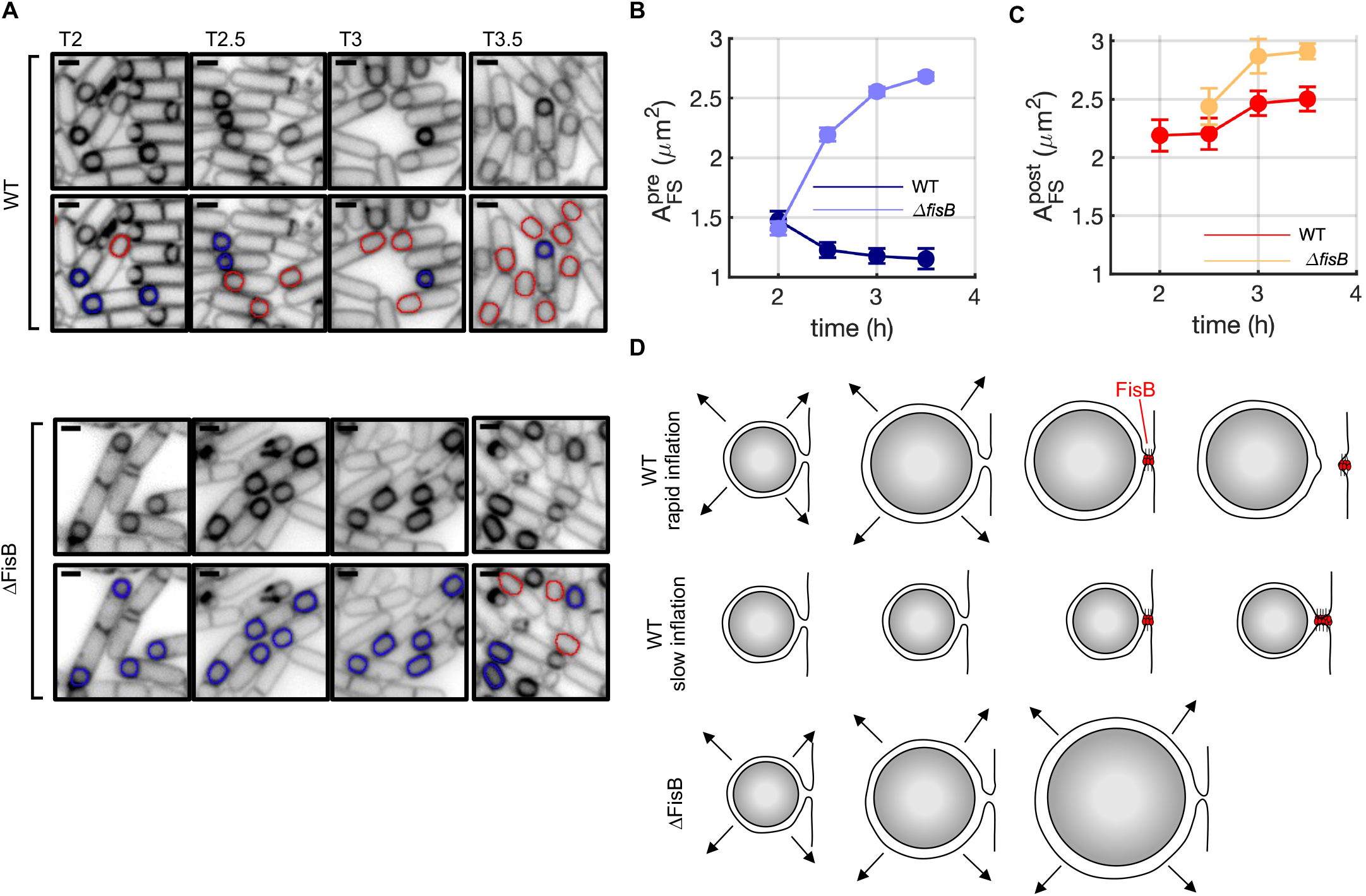
In the absence of FisB, forespores inflate but fission is impaired. **A**. Representative images of wild-type (PY79) or *ΔfisB* cells (BDR1083) at the indicated times after nutrient downshift. Membranes were visualized with TMA-DPH. Lower panels show the FS contours for cells that have (red) or have not (blue) undergone fission. Scale bars represent 1 μm. **B.** The average pre-fission forespore area, Δ£Sθ, grows during sporulation for *ΔfisB* cells, while it decreases for the wild-type strain. **C**. The average post-fission forespore area, 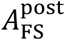, increases faster for *ΔfisB* cells. **D.** Summary of results. FisB catalyzes membrane fission in rapidly growing forespores (top row, “WT rapid inflation”). Slowly inflating forespores are less likely to undergo membrane fission even in the presence of FisB (middle row, “WT slow inflation”). In the absence of FisB, forespores inflate, but fission is rare. See Figure S4 for results with a strain expressing FisB at reduced levels. In B, C, mean±SEM of 3 independent measurements are shown (70 cells per data point).

Sporulating cells expressing GFP-FisB at ~8-fold lower levels than wild-type undergo membrane fission more slowly (Figure S4). In these cells, post-fission forespore areas, 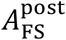, were slightly larger than those expressing GFP-FisB at native levels, and in both cases 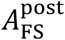 increased over time (Figure S4F). Similar to the *ΔfisB* mutant, the pre-fission forespore area increased as a function of time when GFP-FisB levels were reduced albeit the increase was more modest (Figure S4E).

Overall, these results indicate that forespore inflation occurs independently of FisB, but that forespore inflation is not sufficient to catalyze efficient membrane fission. When FisB is present, the fission process is greatly accelerated by forming an aggregate at the membrane neck to be severed, but only if the forespore is inflating (Figure 5D).

### Modeling suggests increased engulfment membrane tension is the primary driver of FisB-catalyzed membrane fission

Next, we modeled the effect of increased membrane tension on membrane neck geometry, including the roles of hydrodynamic drag on the forespore due the DNA pumping and the osmotic pressure difference between the cytoplasm and the lumen of the neck. We developed a minimal model based on free energy minimization, considering the free energy *F* of an axisymmetric and mirror symmetric membrane neck connecting two membrane sheets, corresponding to the local geometry of the neck formed between the forespore and mother cell membranes (see SI Appendix for details). The energy functional F consists of a term accounting for membrane bending and tension, *F_m_*, another term, *F_p_*, accounting for an osmotic pressure difference Δ*p* = *p*_cyto_ – *p*_peri_ between the cytoplasm and periplasm since the lumen of the neck is continuous with the periplasmic space, and finally a term representing any pulling force *f* on the forespore, e.g. due to DNA translocation, *F_f_*.

We first explored the role of membrane tension, *γ*, on neck geometry. As shown previously (Landajuela et al., 2021) using a minimal model for the neck as a cylinder of radius *r*_cyl_ and length *L*_cyl_, there are two regimes: (1) For a large neck (radius larger than a critical radius *r*_crit_), membrane tension drives the neck to become ever larger. (2) By contrast, for an initial *r*_cyl_ < *r*_crit_, the neck will relax to a finite equilibrium radius set by the balance between membrane tension and membrane bending stiffness. Intuitively, these two regimes follow because as the radius of the cylindrical neck grows, the area of the cylinder grows ~*L*_cyl_ *r*_cyl_, but the areas of the associated flat membranes on the mother cell and forespore *shrink* 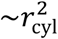. While the former term can dominate for small *r*_cyl_ leading to a local minimum, the latter term eventually dominates for large *r*_cyl_ implying the neck will open. The existence of a local minimum of neck radius is recapitulated by this simplified model in SI Appendix Eqs. 1 and 5. As seen in SI Appendix Fig. 2a, in regime (2) increasing membrane tension *γ* results in a narrowing of the equilibrium neck radius. While to our knowledge there has been no direct measurement of the membrane tension in *B. subtilis*, we find that reasonable values of *γ* = 0.1 – 1 pN nm^-1^ are sufficient to achieve a neck radius < 10 nm (Ojkic et al., 2014). It is likely that cell wall remodeling that drives engulfment (Ojkic et al., 2016) brings the neck radius into regime (2), because a neck forms and a small fraction of cells undergo membrane fission even in the absence of FisB (Figure 1 and Figure 5).

To examine the additional role of osmotic pressure difference between the cytoplasm and the lumen of the neck, we also minimized F at finite Δ*p*. While Δ*p* = 0.1 atm has little effect on the neck radius, Δ*p* = 0.6 atm, which is realistic for *B. subtilis* (Lopez-Garrido et al., 2018; Ojkic et al., 2014) further reduces the minimum neck radius, by e.g. ~1 nm (SI Appendix Fig. 2b and 5), for a fixed value of the membrane tension, *γ* = 0.84 pN nm^-1^. This reduction makes sense, as higher osmotic pressure in the mother cell will tend to compress the volume of the periplasmic space contained within the neck.

By contrast, a pulling force on the forespore not only stretches the neck but also *increases* its equilibrium radius (see SI Appendix Fig. 5). A pulling force of *f* = 50 – 100 pN, around the maximum force that can be applied by one or two SpoIIIE motors translocating DNA into the forespore (Allemand and Maier, 2009; Liu et al., 2018) (see SI Appendix for more details), increases the minimum neck radius significantly compared to a case where there is no pulling force. In particular, the membrane neck radius increases by ~ 4 nm, when *f* = 100 pN, considering *Δp* = 0, *γ* = 0.84 pN nm^-1^ (SI Appendix Fig. 5a). One can understand this effect simply by considering the total force in the neck at the midpoint, which is the product of the circumference of the neck times the membrane tension, *f* = 2*πr*_min_*γ*. So at fixed membrane tension *γ*, an increase of the pulling force *f* automatically implies an increasing *r*_min_. Moreover, an osmotic pressure difference of Δ*p* = 0.6 atm is able to reduce the minimum neck radius by ~1 nm even if the pulling force *f* is not zero, as shown in SI Appendix Fig. 5b, for a fixed value of the membrane tension *γ* = 0.84 pN nm^-1^.

It is likely that the membrane tension in the rapidly expanding forespore and engulfment membranes is higher than in the mother cell membrane. We estimated an upper limit for this membrane tension gradient, by estimating the gradient that would be sufficient to drive a flux of lipids in the neck matching the observed average increase in forespore membrane area (see above and SI Appendix). Perhaps surprisingly, the resulting difference is only ~10”^7^ pN nm^-1^, which is orders of magnitude less than the expected overall membrane tension. Thus an unimpeded flux of lipids between the mother cell and forespore will have negligible effect on the equilibrium results presented above. However, if this flux is strongly impeded, e.g. by accumulation of FisB at the neck, the tension in the engulfment membrane around the forespore would increase further, possibly contributing to membrane fission (see below).

In summary, modeling suggests that the pulling force on the forespore exerted by DNA translocation has a negligible effect on neck geometry. By contrast, a realistically high membrane tension, possibly augmented by an osmotic pressure difference across the membrane, is sufficient to drive the equilibrium neck radius down to ≤ 5-10 nm, with two expected consequences. First, with a small neck radius, FisB can interact in *trans* and accumulate in the neck, facilitating membrane fission (Landajuela et al., 2021). Second, high membrane tension and a small neck radius likely both facilitate membrane fission.

### Membrane fission catalyzed by FisB-lipid friction under high membrane tension

Our results so far suggest that increased membrane tension due to forespore inflation drives FisB-catalyzed membrane fission. Increased membrane tension in the engulfment membrane would provide a driving force for membrane flow from the mother cell to the engulfment membrane through the neck connecting the two compartments. One possibility is that by forming an oligomer at the membrane neck, FisB impedes the movement of lipids that partially support this growth, further increasing the tension in the engulfment membrane, promoting membrane fission. To test this possibility, here we studied lipid flows between membrane compartments in live cells and how lipid mobility is affected by FisB oligomerization.

We first tested if FisB oligomerization on a membrane can impede lipid movement. We incubated FisB ECD with GUVs labeled with fluorescent Cy5-lipids, allowing FisB ECD to form a dense scaffold as in Figure 2. We then probed lipid mobility using FRAP as shown in Figure 6. In the absence of FisB ECD, fluorescence of the bleached region recovered rapidly and nearly completely, indicating unimpeded diffusion, as expected. However, when GUVs were coated with FisB ECD, about half the lipids became immobile (Figure 6C). We conclude that FisB oligomerization severely impedes lipid mobility.

**Figure 6.**
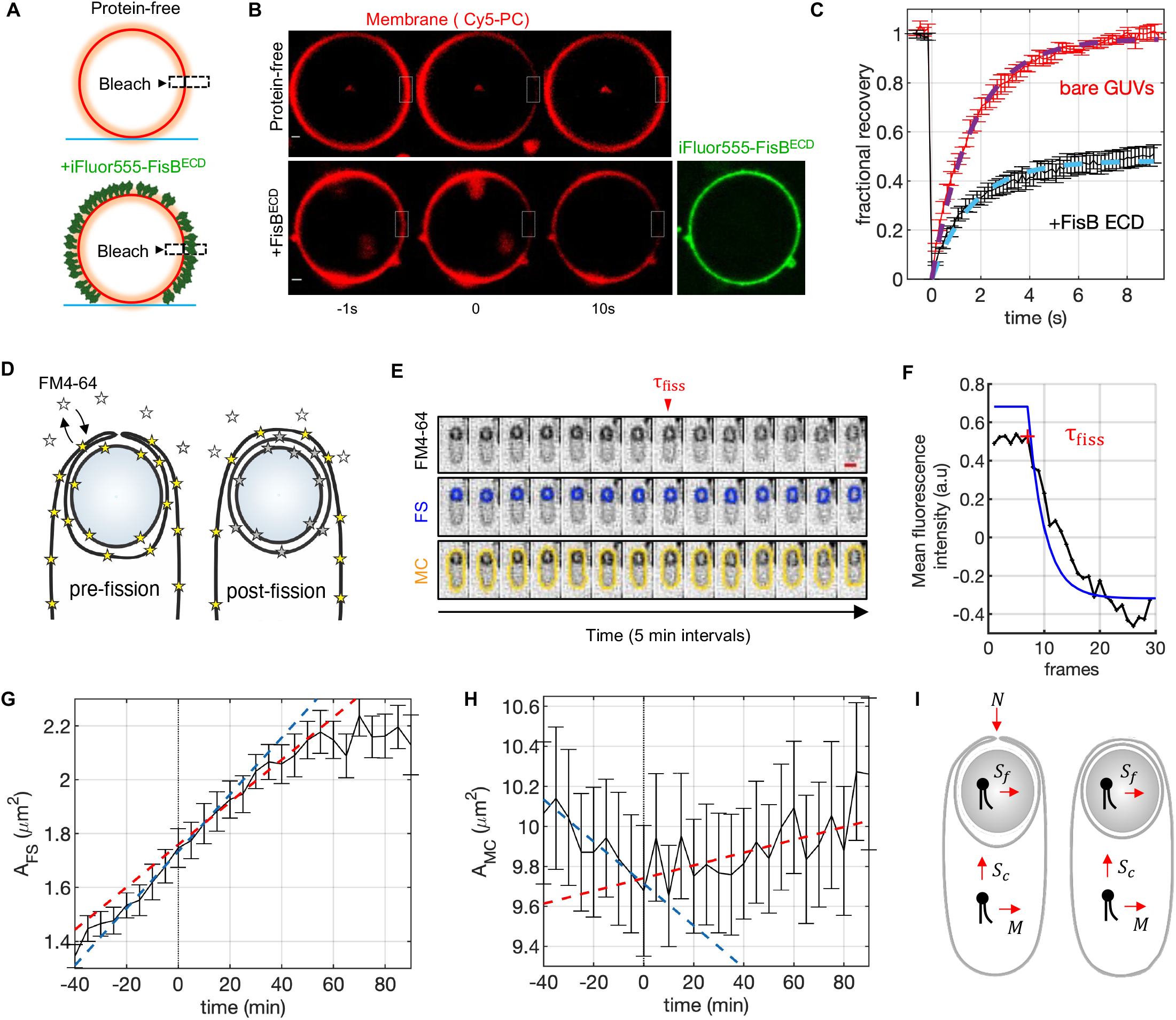
FisB impedes lipid diffusion. **A-C.** Lipid diffusion is impeded by FisB ECD oligomerization on membranes. **A**. Schematic of the experiment. A region of interest (2 *μ*m by 2 *μ*m by ~1 *μ*m in xyz) is bleached and fluorescence recovery is monitored for bare GUVs or GUVs covered by FisB ECD labeled with iFluor555. The schematic is drawn from a side view. Actual bleaching and imaging are performed through the bottom. **B**. Snapshots from time-lapse movies at the indicated times for a representative protein-free (top) or a FisB ECD covered GUV (bottom). Time zero corresponds to the frame right after high-intensity bleach. Scale bar is 1 μm. **C**. Quantification of recovery kinetics. To estimate fractional recovery and an approximate recovery time, the corrected and averaged recovery curves were fitted to *f*(*t*) = /*∞*(1 – exp(-*t/τ*)) where *f*_∞_ and *τ* are the mobile fraction and the recovery timescales, respectively. Best-fit mobile fractions were (with 95% confidence intervals) *f*_∞_ =0.98 (0.97, 0.99) and 0.48 (0.47, 0.49) for bare and FisB ECD coated GUVs, respectively (*R*^2^=0.98-0.99). In both cases *τ* ≈ 1.7 s, implying a lipid diffusivity of a few *μm*^2^/*s*. **D-I.** Membrane flows between membrane compartments before and after membrane fission. **D**. Schematic showing how the timing of membrane fission is detected in time-lapse movies of individual cells. The dye (depicted as a star) becomes highly fluorescent upon insertion into the membrane. The lipid-inserted and free dyes exchange continuously, minimizing bleaching in pre-fission cells. Upon fission, the free dye has no longer access to the inter-membrane space, leading to the onset of an intensity decay. **E.** An example of a single cell, labeled with FM4-64, followed as a function of time (see Figure S5 for more examples). Top row shows a montage of FM4-64 fluorescence images as a function of time. The middle and bottom rows show the detected forespore and mother cell contours respectively, using an active-contour fitting algorithm. The frame preceding fission is indicated in the image (*τ*_flss_). Bar,1 μm. **F**. Plot of the mean FS contour FM4-64 fluorescence (black) as a function of time for the individual cell in E, displaying a relatively stable signal and a sudden decay upon membrane fission. Cross-correlation of the fluorescence signal with a model decay function (blue) was applied to determine the timing of membrane fission *τ*_flss_. **G**. Average of forespore cell membrane surface area of wild-type sporangia as a function of time (n=25). Time axes for individual cells were aligned such that *τ*_fiss_ = 0 min. Error bars represent SEM. The dashed blue and red lines are linear fits to the pre- and fission periods, with slopes 10.6 × 10^/L^ and 7.9 × 10^/L^ μm^2^/min. **H**. Similar to G, but for the mother cell areas. The dashed blue and red lines are linear fits to the pre- and fission periods, with slopes −10.5 × 10^/L^ and +3.2 × 10^/L^ μm^2^/min. **I**. Schematic showing sources and sinks of lipids. Before fission, newly synthesized lipids are inserted into the mother cell and engulfment membranes from the mother cell cytoplasm at rates *M* and *S_c_*, respectively. Lipids synthesized within the forespore are inserted at rate *S_f_*, which likely increases as lipid synthase genes initially located in the mother cell cytopolasm are translocated into the forespore (Pedrido et al., 2013). Lipid flux through the neck (at rate *N*) is only possible before fission.

Next, we tested if forespore inflation leads to membrane movement between the mother cell and engulfment membrane compartments. We followed the kinetics of forespore area growth in individual cells and related the kinetics to membrane fission. Using time-lapse microscopy, we imaged individual sporulating cells labeled with the lipophilic dye FM4-64 (Lopez-Garrido et al., 2018) as shown in Figure 6D,E (see Figure S5 for more examples). While TMA-DPH crosses the cell membrane slowly, FM4-64 does not cross it, which allowed us to detect the time of membrane fission in addition to monitoring membrane areas. The non-fluorescent FM4-64 molecules in the aqueous phase exchange with the membrane-inserted fluorescent ones (Cochilla et al., 1999). Membrane fission isolates the small inter-membrane space from the bath, blocking exchange of bleached dyes with unbleached ones in the bath and accelerating decay of the fluorescence signals arising from the forespore and engulfment membranes. A plot of forespore contour fluorescence as a function of time for individual cells shows a stable signal, which starts decreasing upon membrane fission. We cross-correlated these individual fluorescence profiles with a model decay function to determine the timing of membrane fission (Figure 6F). We defined *t* = 0 as the frame just preceding fission (Figure 6E,F) and aligned all measurements with respect to this time before averaging membrane areas as shown in Figure 6G,H. The rate of forespore area increase slowed gradually after membrane fission (Figure 6G). Remarkably, the mother cell area shrank before and grew after membrane fission (Figure 6H).

Until FisB accumulates at the membrane neck and the membrane undergoes fission, there are two sources of lipids to support the growth of the forespore and engulfment membranes: *de novo* lipid synthesis and lipid flux between the mother cell and engulfment membranes (Figure 6I). After membrane fission, substantial expansion of the forespore and engulfment membranes can only be supported by *de novo* lipid synthesis. Thus, analysis of the rates of area changes can be informative about lipid flux through the neck. We focused on the mother cell area changes for this analysis, because the lipid synthesis rate in the forespore likely increases as a function of time as the lipid synthase genes are gradually translocated into the forespore (Pedrido et al., 2013). The mother cell area changed at rates of −0.011 μm^2^/min (−262 lipids/s) and +0.0032 μm^2^ /min (+76 lipids/s) before and after fission, respectively. Assuming the rates of insertion into the mother cell of newly synthesized lipids pre- and post-fission do not change appreciably, we estimate ~340 lipids/s move through the neck toward the engulfment membrane before FisB accumulation – a significant fraction of the maximal cellular lipid synthesis capacity (see above). Membranes lyse when stretched above only ~1 % in area (Evans et al., 2003). Hence, assuming that the total area of the engulfment and forespore membranes is ~ 5 *μ*m^2^, a deficit of ~300 lipids/s would cause a 1% deficit of the membrane area, i.e. ~ 0.05 *μ*m^2^, within ~ 4 min if the area expanded at fixed number of lipids. Thus, it is possible that FisB oligomerization – by interfering with lipid flux through the neck – leads to a further increase in the engulfment membrane tension and thus facilitates membrane fission.

## DISCUSSION

Bacteria rely on membrane fission for every division cycle and during endospore formation. FisB is the only molecule described so far with a dedicated role in membrane fission in bacteria (Doan et al., 2013; Landajuela et al., 2021), but how FisB catalyzes membrane fission has remained unclear. Here we report that FisB does not remodel membranes directly but forms a stable network on them. Thus, another cellular process must be involved to achieve membrane fission at the end of engulfment. Surprisingly, we found that pumping of the chromosomal DNA into the forespore by the ATPase SpoIIIE – a seemingly unrelated process – is a necessary condition for FisB-dependent membrane fission. DNA pumping into the forespore leads to forespore inflation through increased turgor, stretching the thin layer of peptidoglycan between the forespore and engulfment membranes and smoothing membrane wrinkles (Lopez-Garrido et al., 2018). FisB accumulation at the membrane fission site and forespore inflation are both necessary conditions for efficient membrane fission.

We propose that increased membrane tension in the engulfment membrane due to forespore inflation drives membrane fission (Figure 7). High membrane tension can cause membrane fission through opening of a transient pore (Karatekin et al., 2003), as was shown for fission catalyzed by BAR domain proteins scaffolding a membrane tube under extension (Simunovic et al., 2017). The presence of a protein scaffold on a membrane tube has a dual role: slowing equilibration of membrane tension by impeding membrane flow, and acting as a heterogeneous nucleation point for a membrane defect, lowering the energy barrier for membrane poration (Simunovic et al., 2017). Thus, FisB can similarly have a dual role in membrane fission. First, by oligomerizing at the membrane neck, FisB impedes membrane flow from the mother cell membrane to the engulfment membrane where tension is higher, leading to a further increase in tension, making fission more likely. Second, the FisB oligomer-bare membrane interface could provide a nucleation point for a membrane defect at high tension.

**Figure 7.**
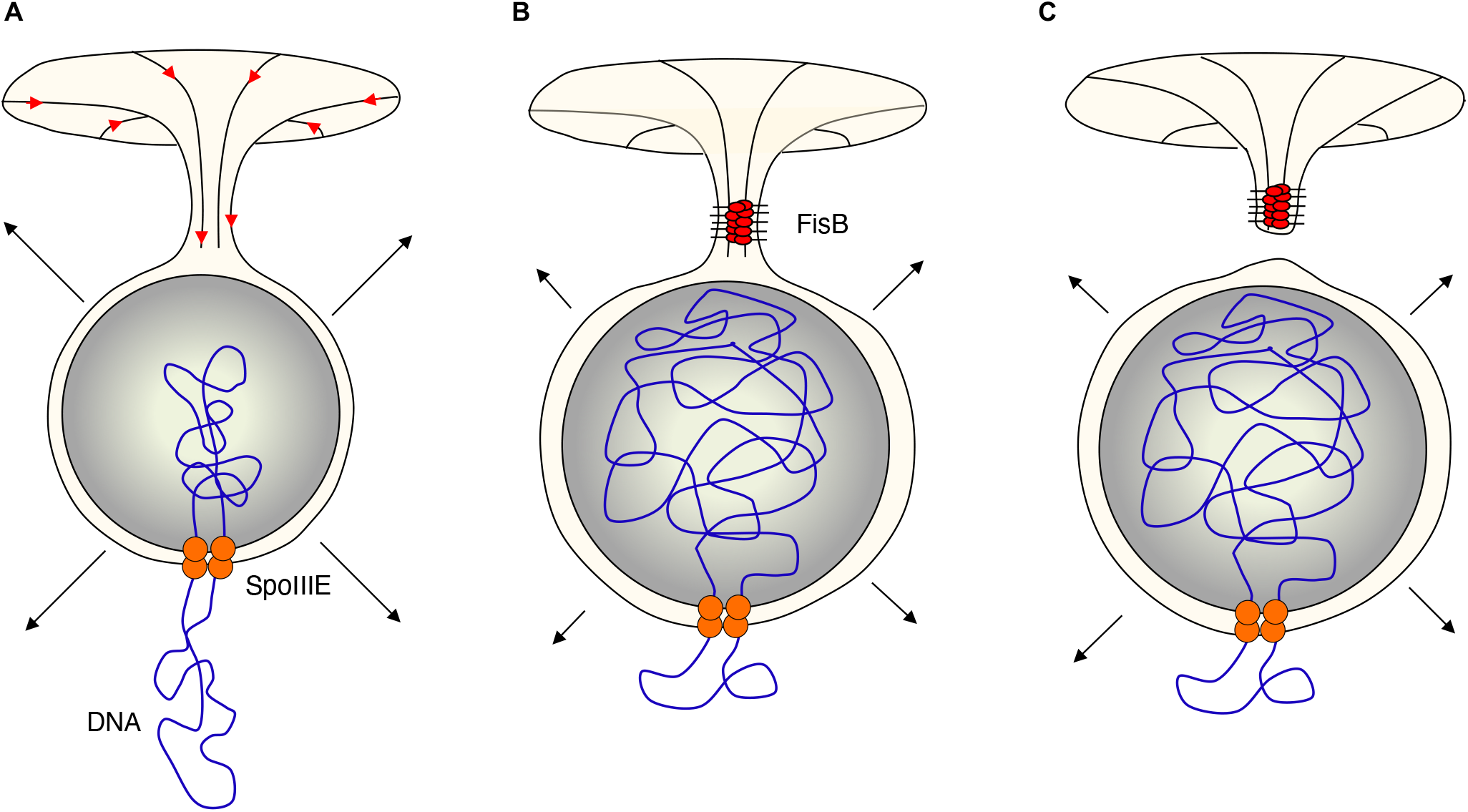
Proposed model of how forespore inflation drives FisB-mediated membrane fission. **A**. DNA translocation by the ATPase SpoIIIE inflates the forespore, stretching the thin peptidoglycan layer between two membranes and smoothing membrane wrinkles (Lopez-Garrido et al., 2018). Increased membrane tension in the engulfment membrane drives lipid flux through the neck between the engulfment and mother cell membranes, partially supplying lipids needed for forespore inflation. **B**. FisB oligomerizes at the membrane neck, impeding membrane flow and causing the engulfment membrane to stretch further. **C**. Increased membrane tension drives membrane scission.

Membrane fission requires energy input. For all biological membrane fission reactions described to date, membrane fission is energized and regulated locally. For the best characterized cases, membrane fission proteins dynamin (Ferguson and De Camilli, 2012) or ESCRT III (Schoneberg et al., 2017) polymerize at the membrane fission site and use the energy liberated by nucleoside triphosphate hydrolysis to reshape membranes locally. By contrast, here we have found that membrane fission during endospore formation in *B. subtilis* is energized indirectly through DNA translocation into the forespore – a process that is required primarily for genetic inheritance and at first sight bears little relation to the membrane fission that occurs at the opposite end of the engulfment membrane. This situation is analogous to secondary transporters using electrochemical gradients set by primary pumps such as the sodium-potassium ATPase to energize secondary transport processes (Boron and Boulpaep, 2017).

Additionally, our results clarify the role of SpoIIIE in membrane fission during endospore formation. It was originally proposed that in addition to DNA translocation, SpoIIIE drives membrane fission because an ATPase deficient mutant resulted in membrane fission defects (Liu et al., 2006; Sharp and Pogliano, 1999). However, SpoIIIE’s role in membrane fission remained unclear because the knock-out or ATPase dead mutants (Figure S3 and (Doan et al., 2013)) cause severe engulfment defects. Since FisB localization and membrane fission occur downstream of engulfment, any engulfment defect would also lead to defects in FisB localization and membrane fission and would not be particularly informative about membrane fission itself. Here we establish that SpoIIIE does indeed play a critical role in membrane fission even in the absence of engulfment defects, but this role is related to forespore inflation.

The membrane fission mechanism we have described here provides an elegant example of how bacteria can use energy very efficiently under starvation conditions. DNA translocation into the forespore is an energy-consuming process that cannot be bypassed, but the byproduct of this process, namely increased turgor in the forespore, can be harnessed to energize secondary processes. It will be interesting to see if processes other than membrane fission are energized by mechanical energy stored in the inflated forespore.

## ACKNOWLEDGEMENTS

We thank all members of the Karatekin laboratory for fruitful discussions and Jeorg Nikolaus (director of the Yale West Campus Imaging Core) for help with imaging. This work was supported by National Institute of General Medical Sciences and National Institute of Neurological Disorders and Stroke of the National Institutes of Health (NIH) under award numbers R01GM114513 and R01NS113236 (to EK). The content is solely the responsibility of the authors and does not necessarily represent the official views of the National Institutes of Health. This work was supported in part by the National Science Foundation, through the Center for the Physics of Biological Function (PHY-1734030) and was performed in part at the Aspen Center for Physics, which is supported by National Science Foundation grant PHY-1607611. AMC also acknowledges support from the Princeton Center for Theoretical Science and the Human Frontier Science Program through the grant LT000035/2021-C. We gratefully acknowledge a Yale University Predoctoral Fellowship to MB.

## AUTHOR CONTRIBUTIONS

EK, MB, AL, and DZR conceived the study. MB carried out most of the experiments with purified proteins and artificial membranes, AL carried out live cell imaging and analysis. AL and CGP performed the lipid mobility measurements. MB, EK wrote matlab scripts for image analysis, AL, MB analysed images. AMC and NSW developed the mathematical model, DZR, NSW and EK supervised the work. CDAR and TD provided reagents and intellectual input. EK wrote the manuscript, with input from all authors.

## STAR★Methods

### Key Resources Table

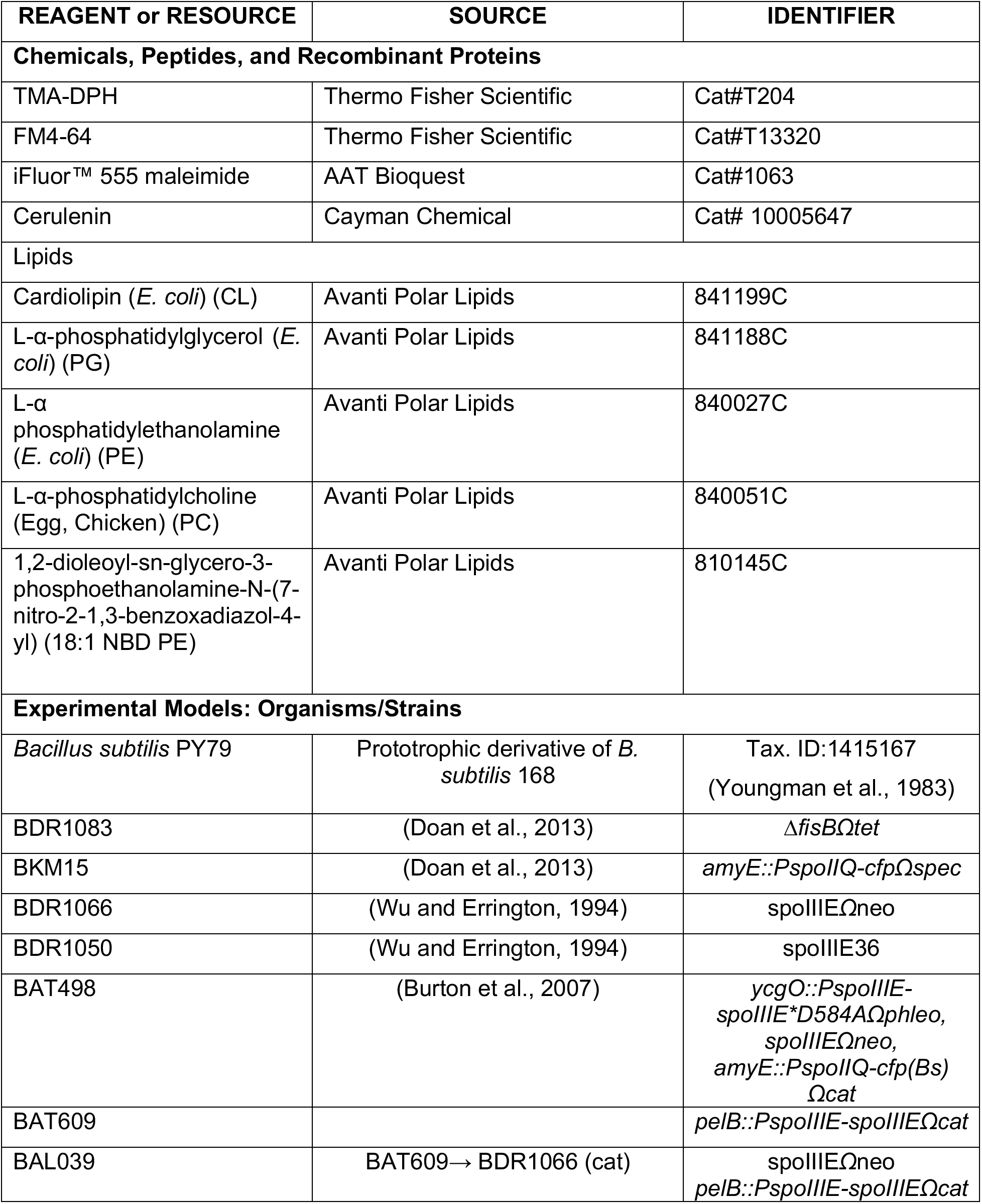
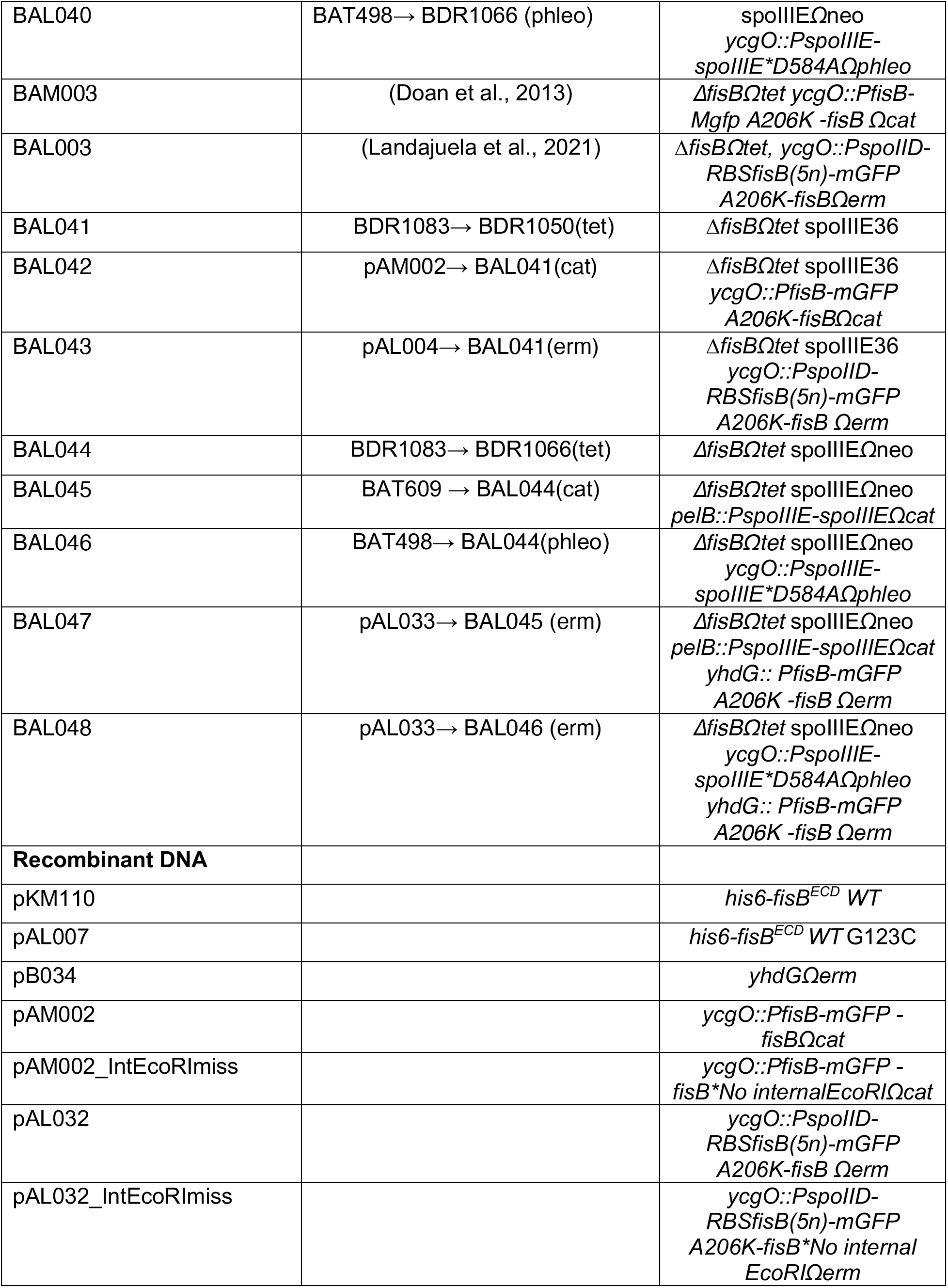
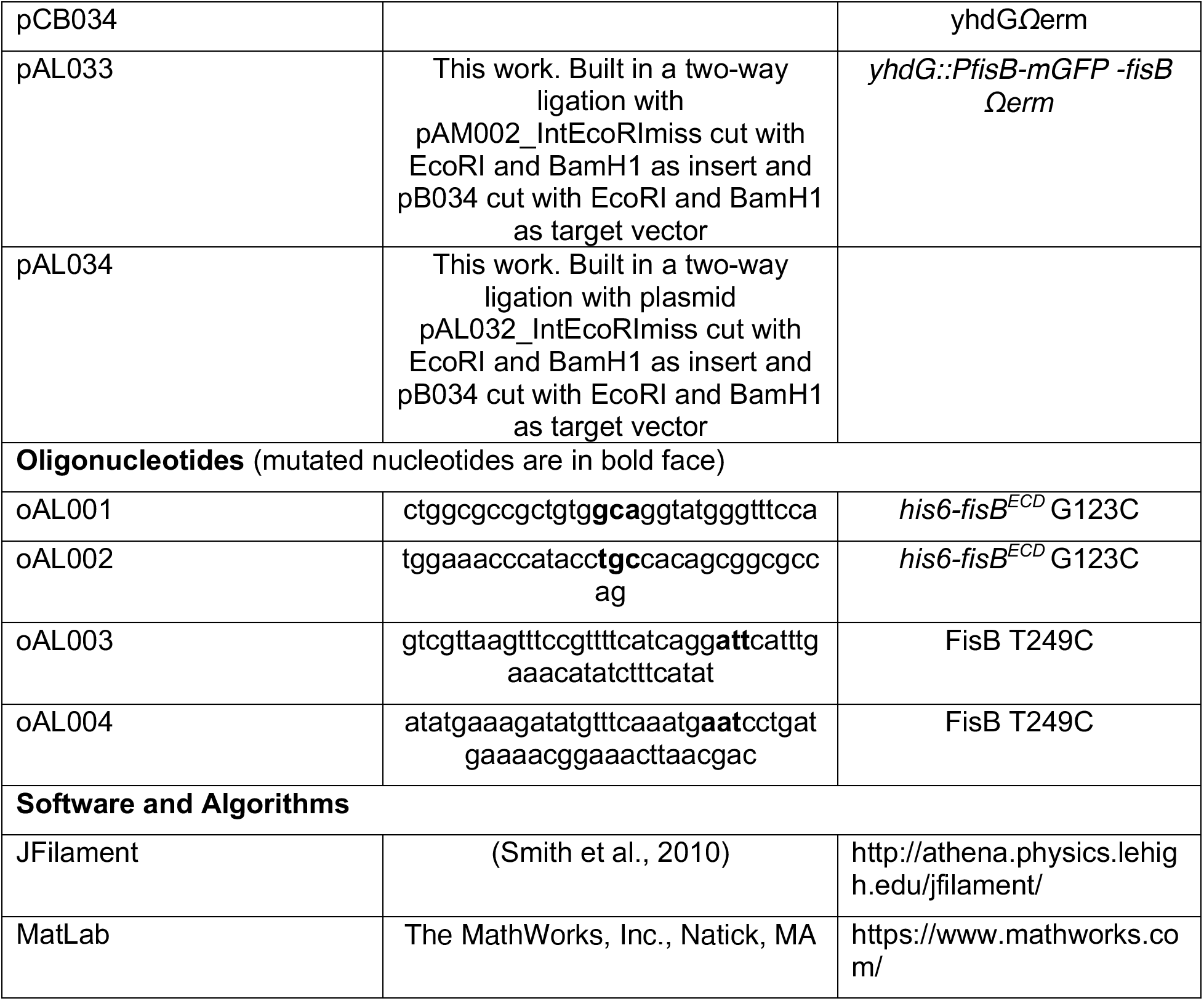

### Contact for Reagent and Resource Sharing

Further information and requests for resources and reagents should be directed to and will be fulfilled by the Lead Contact, Erdem Karatekin

## Method details

### Experimental Procedures

#### General *B. subtilis* methods

*B. subtilis* strains were derived from the prototrophic strain PY79 (Youngman et al., 1983). Sporulation was induced in liquid medium at 37°C by resuspension according to the method of Sterlini-Mandelstam (Sterlini and Mandelstam, 1969). *B*. *subtilis* strains were constructed using plasmidic or genomic DNA and a 1-step competence method. Site directed mutagenesis was performed using Agilent’s Quick-change Lightning kit following manufacturer’s instructions and mutations were confirmed by sequencing. The list of strains and plasmids can be found in the Star Methods Key Resources Table.

#### Fluorescence microscopy from batch cultures

500 μl samples at indicated times during sporulation were taken and concentrated ~50-fold by centrifugation (3300xg for 30 s). Membranes were stained with 1-(4-trimethylammoniumphenyl)-6-phenyl-1,3,5-hexatriene *p*-toluenesulfonate (TMA-DPH; Molecular Probes) at a final concentration of 100 μM and the cells were then immobilized on a 2% agarose pad made with sporulation buffer covered with a no.1.5 coverslip, after which the cells were immediately imaged live. In most cases fluorescence microscopy was performed using a Nikon Ti microscope equipped with a ×100 Plan Apo 1.45 NA phase-contrast oil objective, an Orca-Flash4.0 V2 CMOS camera (Hamamatsu Photonics, Shizuoka,Japan) and a Spectra X light engine (Lumencor, Beaverton, OR, USA), all controlled by Nikon Elements software (Nikon Corp. Tokyo, Japan). Excitation of TMA-DPH was achieved using the 395/25 nm band of the SpectraX system and Chroma’s ET 49000 single band filter set the DAPI filter set. (λ_ex_=395/25 nm; λ_em_=431/28 nm). Excitation light intensity was set to 50% and exposure times were 200 ms.

In experiments shown in Figure 6 and S3, cells were prepared following the same procedure but visualized on a Leica DMi8 Wide-field Inverted Microscope equipped with a HC PL APO 100× DIC objective (NA=1.40), and an Andor iXon Ultra 888 EMCCD camera (Andor Technology Ltd., Belfast, Northern Ireland) and a Spectra X light engine (Lumencor, OR,USA) controlled by the LAS X program (Leica Microsystems, Wetzlar, Germany). Excitation of TMA-DPH was achieved using the 395/25 nm band of the Spectra X system and the Leica DAPI filter set (λ_ex_=395/25 nm; λ_em_=460/50 nm). Excitation light intensity was set to 50% and exposure times were 300 ms..

Images were processed using the ImageJ software. Contours generated using JFilament were coloured using Photoshop.

#### Time-lapse fluorescence microscopy

Cells were visualized on a Leica DMi8 Wide-field Inverted Microscope equipped with a HC PL APO 100× DIC objective (NA=1.40) and an iXon Ultra 888 EMCCD Camera from Andor Technology. Sporulation was induced at 37°C. To visualize the membranes, 0.5 μg/ml FM4-64 was added to the culture ~1 hours after sporulation induction and incubation continued for another hour. Then a 10 μl sample was taken and transferred to an agarose pad prepared as described (Lopez-Garrido et al., 2018). Pictures were taken in an environmental chamber at 37°C every 5 min for at least 2 hours. Excitation of FM4-64 was achieved using the 575/25 nm band of the SpectraX system and a custom FM4-64 filter set (λ_ex_=395/25 nm; λ_em_>610nm). Excitation light intensity was set to 5% to minimize phototoxicity and exposure times were 100 ms. For presentation purposes, sporangia were aligned vertically (with forespore on top) by rotating them using ImageJ. Contours generated with JFilament were colored using Photoshop.

#### Expression, purification, and labeling of recombinant proteins

FisB ECD and mutants were purified as described in (Landajuela et al., 2021). Briefly, Hisβ-FisB ECD was expressed in *E. coli* BL21 (DE3) (New England Biolabs, Ipswich,MA,USA) and purified using HisPur™ Ni-NTA Resin (Thermo-Fisher Scientific, Waltham, MA, USA). Protein expression was induced with 1 mM IPTG at OD_600_ = 0.6 overnight at 16°C. Cells were harvested by centrifugation and the pellet was resuspended in Lysis Buffer (20 mM HEPES, 500 mM NaCl, 0.5 mM TCEP, 20 mM imidazole, 2% glycerol, 20 mM MgCl2) and flash frozen in liquid nitrogen. Pellets were thawed on ice and cells were lysed by 5 passes through a high-pressure homogenizer (Avestin EmulsiFlex-C3, Ottawa, Canada). The lysate was spun down at 100,000 g and the soluble fraction was incubated with HisPur™ Ni-NTA Resin for 2.5 h at 4°C while rotating. The bound protein was washed sequentially with Lysis Buffer, Lysis Buffer containing 50 mM and finally 100 mM Imidazole. The protein was eluted in Elution Buffer (20 mM HEPES, 500 mM NaCl, 0.5 mM TCEP, 200 mM Imidazole, 2% glycerol, 20 mM MgCl2). The protein was concentrated using a Vivaspin® with a 10 kDa molecular weight cutoff and the concentration determined by Bradford protein assay. His6-FisB ECD G123C was labeled with iFluor555™ maleimide dye (AAT Bioquest, Sunnyvale, CA, USA) by adding a 20x molar excess of dye to the protein and the reaction was incubated overnight at 4°C. Free dye was removed using a HiPrep 20/10 (GE, Chicago, IL, USA) column.

#### Giant Unilamellar Vesicle (GUV) preparation

GUVs were formed by electroformation (Stockl et al., 2010). Briefly, chloroform-dissolved lipids were mixed in a glass tube at desired ratios and spotted on two indium tin oxide (ITO) coated glass slides. Organic solvent was removed by placing the lipid films in a vacuum desiccator for at least 2 h. A short strip of copper conductive tape was attached to each ITO slide which are then separated by a PTFE spacer and held together with binder clips. The chamber was filled with 500 μl Swelling Buffer (SweBu, 1 mM HEPES, 0.25 M sucrose, 1 mM DTT) and sealed with Critoseal (McCormick Scientific LLC, Saint Louis, MO, USA). GUVs were formed by applying a sinusoidal voltage of 10 Hz and an amplitude of 1.8 V for at least 2 h at room temperature. GUV membranes were labeled by including 1 mole % 1,2-dioleoyl-sn-glycero-3-phosphoethanolamine-N-(7-nitro-2-1,3-benzoxadiazol-4-yl) (NBD-PE) or 1,1’-dioctadecyl-3,3,3’,3’-tetramethylindodicarbocyanine perchlorate (DiD) in the lipid composition. In Figure 2, “100% PC” and “30% CL” membranes are composed of (all mole %) 99% phosphatidylcholine (PC), 1% NBD-PE, and 30% *E. coli* cardiolipin (CL), 69% eggPC, 1% NBD-PE, respectively. Deflated GUVs were composed of (all mole %) 25% *E. coli* phosphatidylethanolamine (PE), 5% *E. coli* CL, 50% *E. coli* phosphatidylglycerol (PG), 19% eggPC and 1% DiD or NBD-PE (BS mix). All lipids were purchased from Avanti Polar Lipids (Alabaster, AL), except for DiD which was from Thermo Fisher (Waltham, MA).

GUVs were imaged using a Nikon Eclipse TE2000-E microscope. Prior to adding the GUVs, the imaging chamber was filled with a 5 mg/ml β-Casein (Sigma, Saint Louis, MI, USA) solution to prevent attachment of the GUVs to the glass coverslip. The β-Casein solution was removed and replaced with RB-EDTA.

#### Fluorescence recovery after photobleaching (FRAP)

##### Protein mobility

FRAP measurements were conducted using a Leica SP8 inverted microscope in the same open imaging chamber. As described above, GUVs were diluted in RB-EDTA (25 mM HEPES at pH 7.4, 850140 mM KCl, 1 mM EDTA, 0.2 mM tris(2-carboxyethyl) phosphine) and incubated with 1 μM FisB ECD for 2-3h. A rectangular area was chosen to bleach as indicated in Figure 2C. Five images (laser power 0.7%) were recorded before bleaching, 10 during bleaching (laser power 100%) and 60 after bleaching (laser power 0.7%).

The mean GUV intensity was determined using ImageJ. A segmented line (10 pixels wide) was drawn to manually follow the GUV membrane. The line was smoothed using ‘fit spine’ and the mean pixel intensity was calculated. The background was determined as the mean pixel value of a 20 x 20-pixel box (close to but outside the GUVs) and subtracted from the mean pixel intensity of the GUVs.

##### Lipid mobility

To monitor lipid mobility in the presence or absence of FisB ECD, GUVs (E.ColiPE:E.ColiCL:E.ColiPG:EggPC:Cy5-PC = 25:5:50:19:1) were prepared as described above and deposited on top of a 0.1% poly-lysine treated glass-bottom dish. After incubation for 30 min at room temperature, the poly-lysine solution was removed and replaced with RB-EDTA. For some samples, 1 μM iFluor555-FisB ECD was added and incubated with the GUVs for 1.5 h. Unbound FisB-ECD was washed with RB-EDTA before imaging. Photobleaching was performed using a Leica SP8 inverted confocal microscope by scanning the 633 nm and 649 nm beams operating at 100% laser power over a rectangular region of interest (ROI, 2 μm by 2 μm). The focal plane was set to mid-GUV height. Note that the bleached membrane area is the cross-section of the ROI set in the imaging plane and the optical thickness (~1 μm), i.e. ~ 2 μm^2^. Five frames were acquired at low (1%) laser power for normalization of the fluorescence signal before bleaching the ROI for 1.53 s and recovery was monitored for 60 frames every 154 ms at low laser power (1%). Image size was 128 x 128 pixels. The background was determined with the same ROI area outside the vesicle using ImageJ and subtracted from the signal. The mean pixel intensity of the entire GUV was used to correct for photobleaching during low intensity excitation read-out after rescaling to account for polarization effects. Because the unbleached lipid reservoir is finite, the mean pixel intensity of the entire GUV decreases during the high-intensity bleach. The GUV contour intensity just after bleaching was used as the maximum possible recovery value when calculating the fractional recovery of the ROI fluorescence. To estimate fractional recovery and an approximate recovery time, the corrected and averaged recovery curves were fitted to *f*(*t*) = *f*_∞_(1 – exp(-*t*/*τ*)) where *f*_∞_ and *τ* are the fractional recovery and the recovery timescales, respectively.

### Image analysis

#### Determination of forespore and mother membrane surface area

Using the ImageJ plugin Filament 2D (Smith et al., 2010), “snakes” were fitted to forespores and mother cells (Figure S1A). Snakes were fitted to forespores using either the TMA-DPH membrane stain (using ridges) or a soluble CFP marker (using gradients) expressed in the forespores. Snakes were fitted to mother cells using the corresponding phase contrast image using gradients. In all cases the snakes were fitted as contours. Fitting parameters were as follows: α = 100, β = 100, γ = 800, weight=0.5, stretch force=100, deform iterations=50, point spacing=0.5, image smoothing=1.01.

To determine the surface area of forespores and mother cells, the snakes were then further analyzed with MATLAB. First, an ellipse was fitted to the snakes to determine the symmetry axis of the forespore or mother cell (Figure S1B). Half of the snake is then rotated around the symmetry axis to create a surface of revolution from which the membrane surface area for the forespore or mother cell was determined (Figure S1C).

For the analyses shown in Figure 5I, FisB foci were semi-automatically selected using SpeckleTrackerJ (Smith et al., 2011). For each spot, the sum of pixel values in a 6 pixels × 6 pixels (0.5 μm×0.5 μm) box around the center of the spot were calculated. For each corresponding cell, the same operation was performed at a membrane area where no clusters were present and subtracted from the individual FisB cluster intensity.

#### Determination of mean fluorescence intensity around the forespore

We used ImageJ plugin Filament 2D to fit snakes around forespores (see above) to define forespore contours. Mean contour intensity was calculated using Matlab, using snake coordinates from Filament 2D dilated to 4 pixels used as a mask.

#### Modeling

Please see SI Appendix for details.

## SUPPLEMENTARY FIGURE LEGENDS

**Figure S1.**
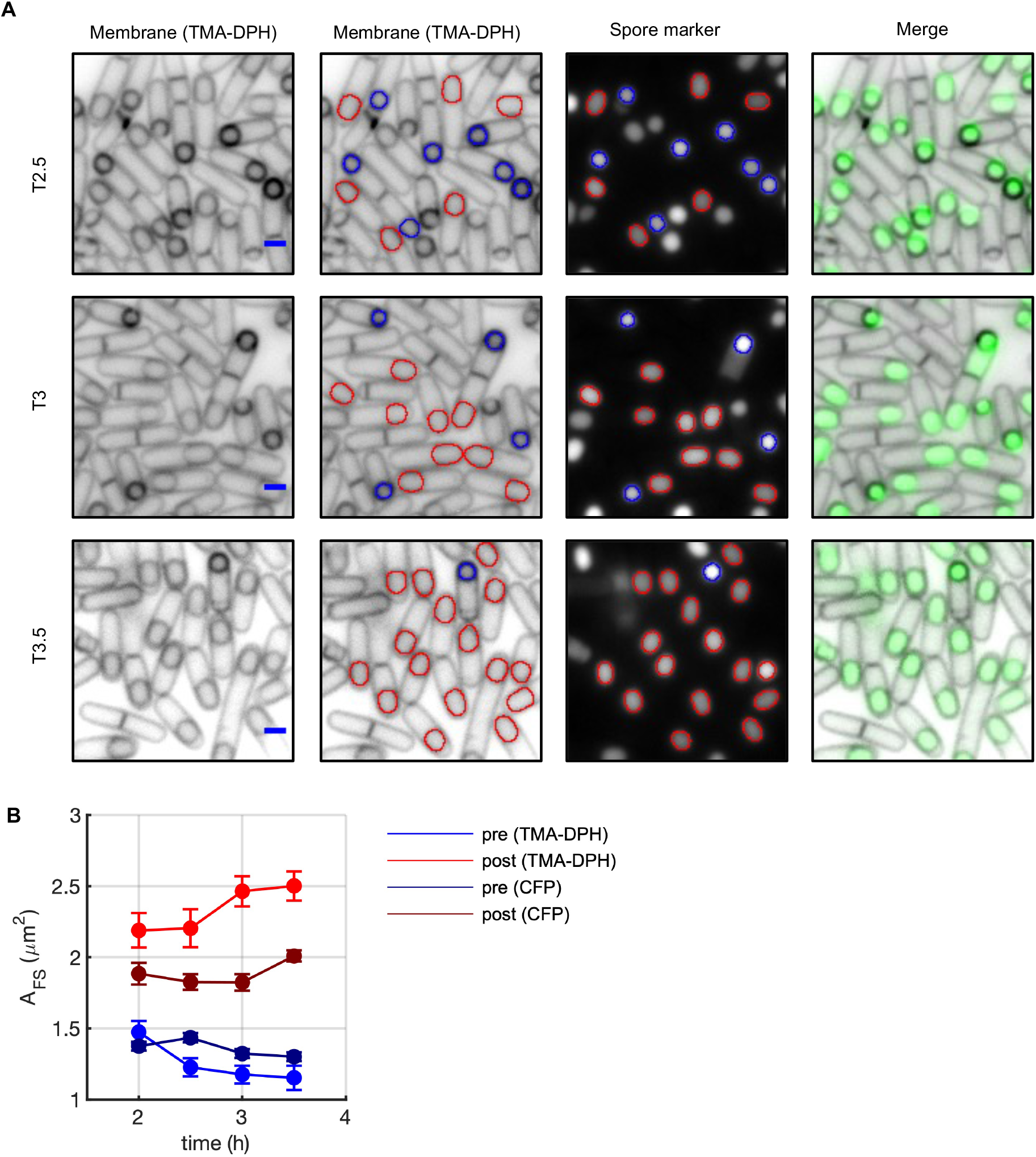
Forespore area changes can also be detected using a soluble marker. Related to Figure 3. **A.** Membrane fission was assessed by fluorescent microscopy during a sporulation time course in wild type (BKM015) cells containing a fluorescent forespore reporter (*P_spoIIQ_-cfp*). Membranes were visualized with the fluorescent dye TMA-DPH. Forespore contours in pre- and post-fission cells are indicated with blue or red contours, respectively. Time (in hours) after the initiation of sporulation is indicated as T2.5, T.3, etc.. Bar, 1 μm. **B.** Average post-fission forespore areas (*A*_FS_) detected using either marker grow while average pre-fission FS areas shrink as a function of time into sporulation. Error bars represent SEM from three independent experiments (70 cells were analyzed per point).

**Figure S2.**
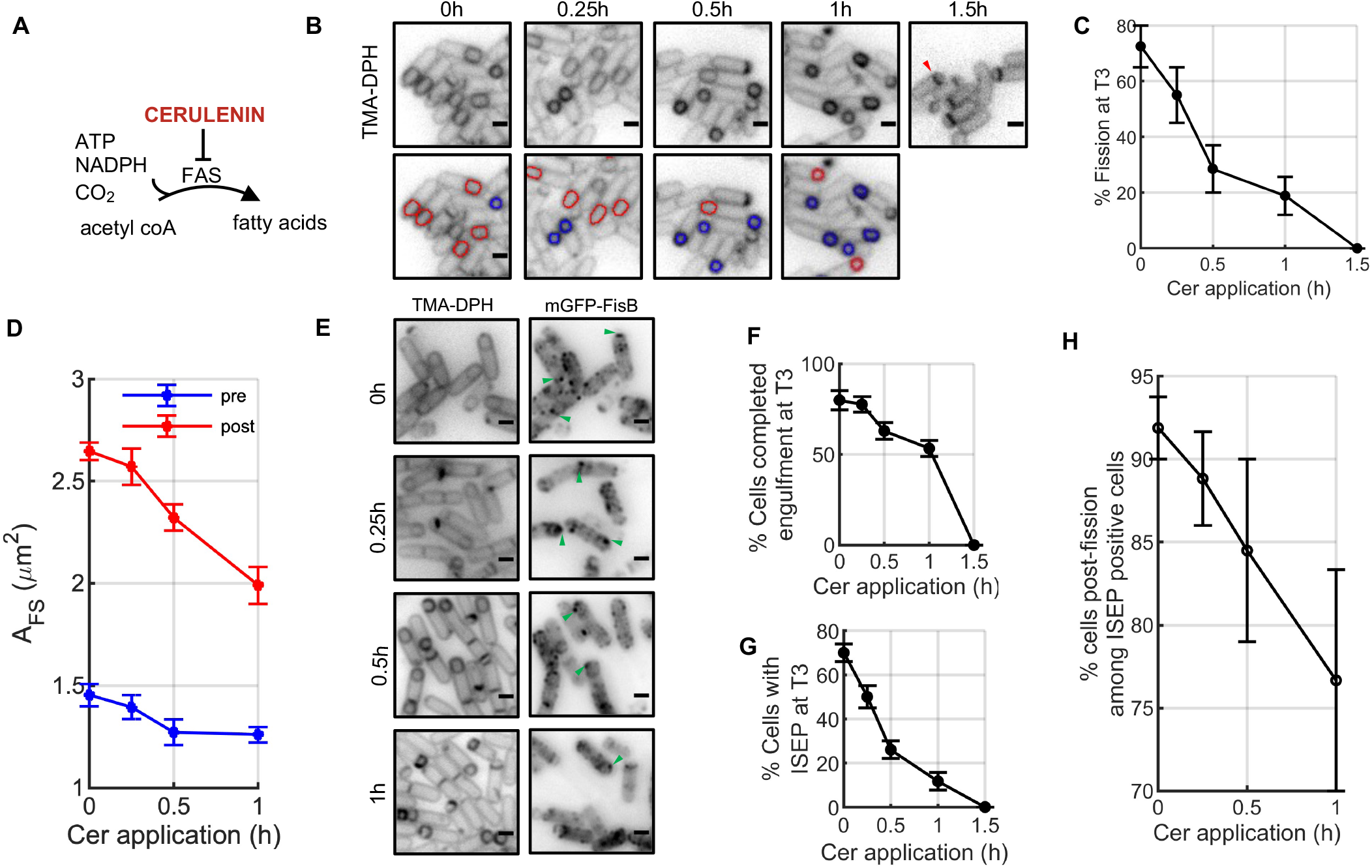
Blocking lipid synthesis inhibits forespore inflation and membrane fission. Related to Figure 4. **A.** Cerulenin (Cer) inhibits *de novo* synthesis of neutral lipids and phospholipids by inhibiting *de novo* fatty acid biosynthesis. **B.** Representative images showing cells at *t* = 3 h after addition of 10 μg/ml cerulenin at indicated timepoints after nutrient downshift. Membranes were visualized with TMA-DPH. A cell that is blocked at the asymmetric division stage is highlighted (red arrowhead). Scale bars = 1 μm**. C.** Percentage of cells (wild-type strain, PY79) that have undergone membrane fission at *t* = 3 h after addition of cerulenin at the indicated timepoints. More than 300 cells were analyzed for each time point. **D**. Quantification of forespore membrane area for cells that have (red) or have not (blue) undergone membrane fission at *t* = 3 h after cerulenin addition at the indicated times after nutrient downshift (n=25-70 cells per data point). **E.** Fractional area occupied by the FS in pre- (blue) and post-fission (red) cells at t = 3 h. Cerulenin was added at the indicated times after nutrient downshift (n=25-70 cells per data point). **F**. Representative images showing cells (strain BAM003) at t=3 hr after the nutrient downshift, with Cer application for the indicated durations. Examples of cells with a discrete mGFP-FisB focus at the cell pole are highlighted with green arrowheads. Scale bars represent 1 μm. **G.** Engulfment is perturbed to a greater extent when lipid synthesis is inhibited earlier by cerulenin addition (>300 cells per data point). **H**. Percentage of cells with FisB accumulation (ISEP formation) at the membrane fission site, for cells that have visually completed engulment. **I**. The fraction of post-fission cells among cells with correct FisB localization (ISEP formation) as a function of increasing cerulenin application time (n=90-200 cells were analyzed per point). Means±SEM of 3 independent experiments are shown for panels C-E and G-I.

**Figure S3.**
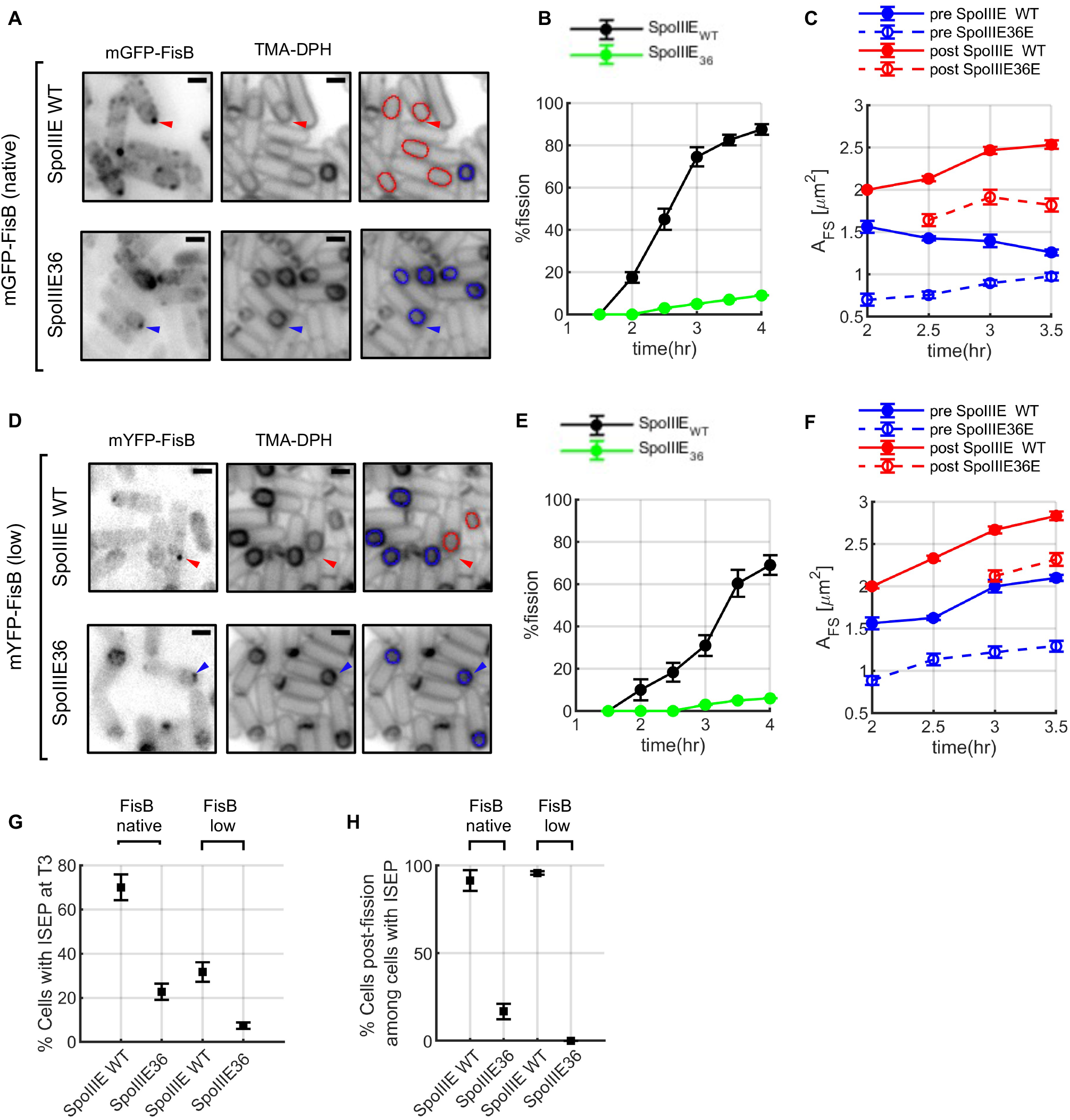
Blocking forespore inflation inhibits membrane fission. Related to Figure 4. **A.** Engulfment and FisB localization in cells expressing wild-type (WT) SpoIIIE or ATP-ase deficient SpoIIIE36. Membranes were labeled with TMA-DPH and FisB was visualized using mGFP-FisB expressed at native levels on a *DfisB* background (Landajuela et al., 2021). Fluorescence microscopy images were acquired 3 h after nutrient downshift that initiated sporulation. Cells that have visually completed engulfment and had accumulated FisB a the membrane fission site are indicated by arrowheads. On the images on the right, detected FS contours are overlaid in blue and red for pre- and post-fission cells. **B**. Percentage of cells that underwent membrane fission as a function of time into sporulation, for WT and SpoIIIE36 cells. (For each data point >300 cells were analyzed). **C**. FS areas as a function of time after nutrient downshift, for pre- (blue) and post-fission (red) cells, expressing either WT (solid lines) or ATPase deficient SpoIIIE (SpoIIIE36). (n=25-70 cells per data point) **D-F**. As in A-C, but using cells expressing mGFP-FisB at ~8-fold lower levels (Landajuela et al., 2021). **G**. FisB localization is affected when DNA translocation is completely blocked. Percentage of cells displaying an intense spot at the engulfment pole (ISEP) for cells expressing SpoIIIE^WT^ or SpoIIIE36 on a background with either native or reduced mGFP-FisB levels. Scale bars = 1 μm. **H.** The fraction of post-fission cells among cells with correct FisB localization (ISEP formation). (n=25-70 cells were analyzed per data point). In panels B,C, E-H, mean±SEM of three independent biological replicates are plotted for every point.

**Figure S4.**
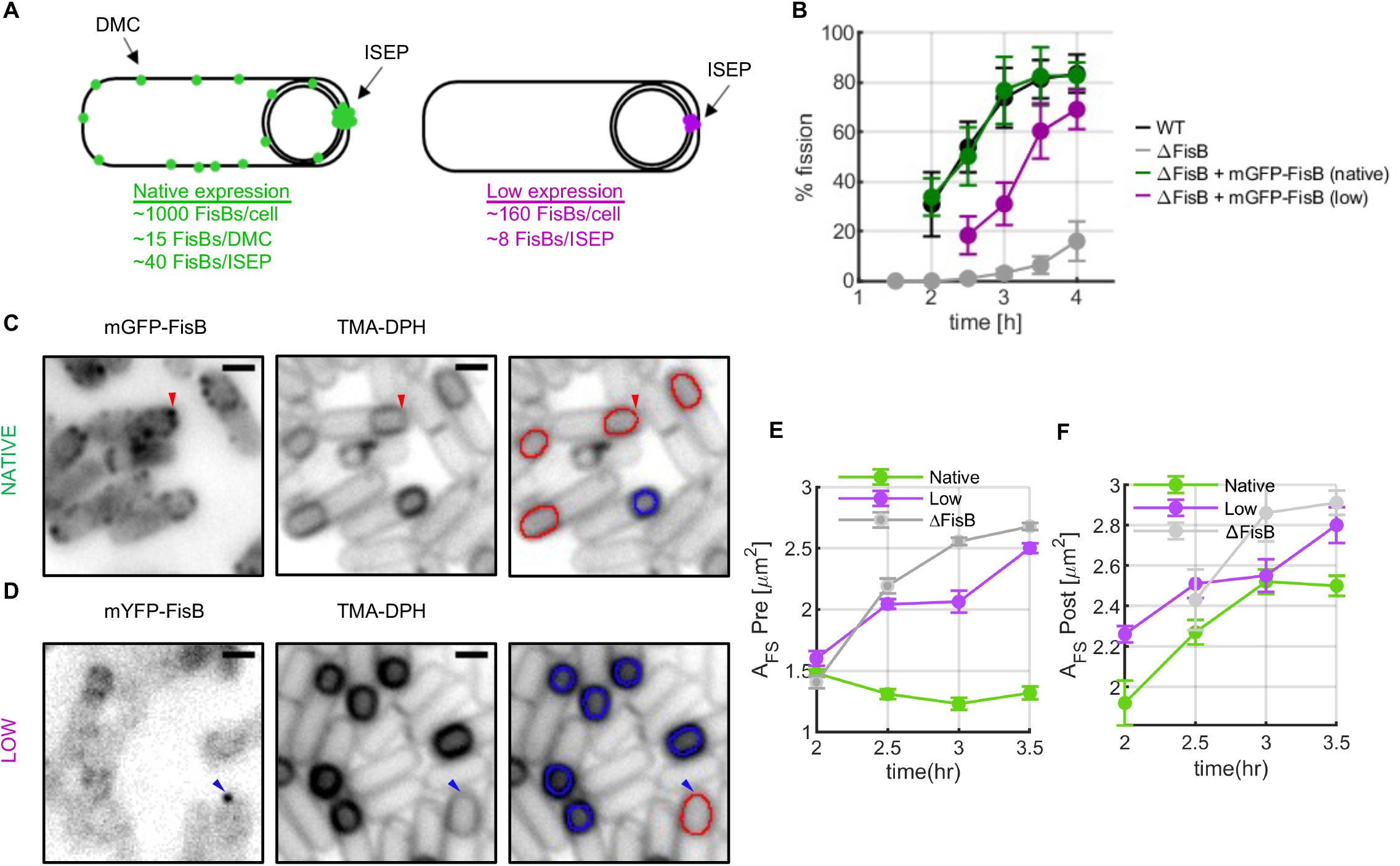
Pre-fission forespores inflate in a low expression FisB strain in which membrane fission is delayed. Related to Figure 5. **A**. Summary of FisB copy number quantification under native or low expression levels (Landajuela et al., 2021). Right. Under native expression, on average, there are ~1,000 FisB molecules per cell at t=3 h. Each DMC contains ~12 FisB molecules, while the ISEP contains ~40 FisB copies. In the low-expression strain all the numbers are scaled down ~7-8-fold. **B.** Time course of membrane fission for wild-type cells, *ΔfisB* cells, or *ΔfisB* cells complemented with mGFP-FisB expressed at native (BAM003) or low levels (BAL003). (More than 300 cells were analyzed per data point.) **C.** Fluorescence microscopy images of cells expressing mGFP-FisB at the native level (BAM003) at 3 h into sporulation. Membranes were visualized with TMA-DPH. Examples of a sporulating cell with a discrete mGFP-FisB focus at the cell pole (intense spot at engulfment pole, ISEP) are highlighted with a red arrowhead. In the panels on the right red contours indicate forespores that have undergone fission whereas blue contours indicate forespores that have not yet undergone membrane fission (70 cells per data point). Scale bars represent is 1 μm. **D.** Similar to C but using a strain (BAL003) that expresses mGFP-FisB at lower levels in a *ΔfisB* background. **E, F.** Under mGFP-FisB expression at low levels, both the average pre- (E) and post-fission (F) forespore areas (*A_FS_*) grow, in a manner qualitatively similar to cells lacking FisB altogether (ΔFisB). By contrast, for cells expressing mGFP-FisB the native level, the average pre- and post-fission FS areas decrease and increase as a function of time into sporulation, respectively (70 cells were analyzed per data point). Data for *ΔfisB* cells is copied from Figure 6 for comparison. In panels B, E, F, mean±SEM of three independent replicates are plotted.

**Figure S5.**
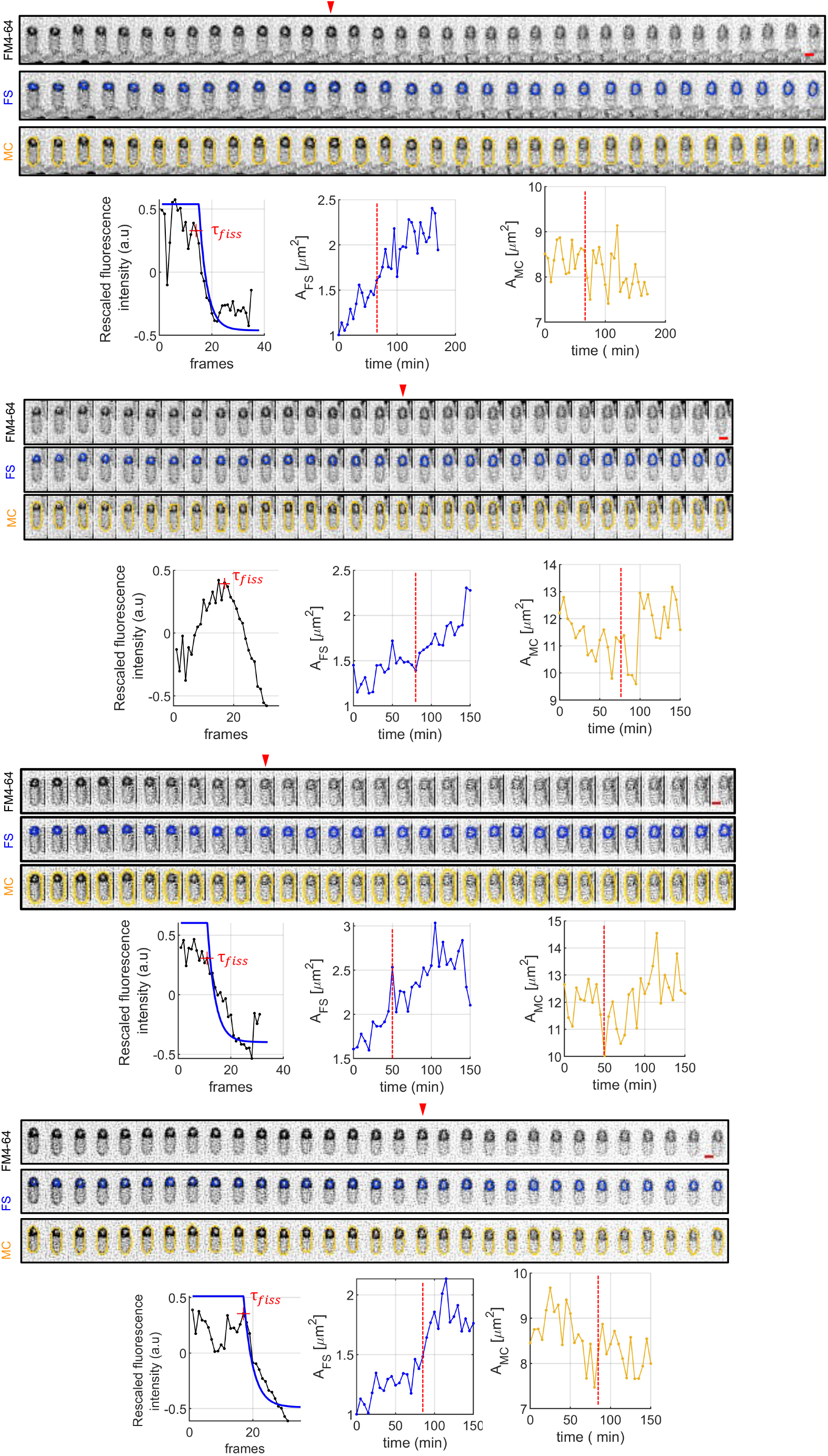
Examples of forespore area growth and membrane fission detected at the single cell level. Related to Figure 6. Each top row of a triplet shows a unique cell followed as a function of time in time-lapse fluorescence microscopy. Cell membranes were labeled with FM4-64. Images are shown with an inverted look-up table. Frames were acquired every 5 min. The middle and bottom rows of every triplet show the overlaid contour of the FS or the MC membrane, respectively. The intensity of the FS membrane starts decreasing upon membrane fission, because bleached FM4-64 molecules are confined and can no longer exchange with the unlabeled dyes in the bath after membrane fission. For each cell, the mean FS contour intensity and the FS area are shown on the right. The time of membrane fission (*τ_fiss_*) estimated from the contour intensity time profile is indicated on the FS area plots.

## SI Appendix Theoretical Modeling

In previous work [1] we showed that a neck of radius ≤ 5 — 10 nm in the region where the engulfment membrane meets the rest of the mother cell membrane (see sketch in Fig. 1a) is necessary for FisB to accumulate. This accumulation in turn favors membrane fission, as shown in the current main text. In this appendix, first, we develop a minimal model for the neck based on free-energy minimization. This minimal model assumes a highly simplified geometry of the membrane neck, as depicted in Fig. 1(b), that accounts for membrane tension, bending, and an osmotic pressure difference, 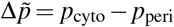, between the cytoplasm and periplasm since the lumen of the neck is continuous with the periplasmic space. Using this model, we explore if an increase of membrane tension in the neck, e.g. due to forespore inflation, along with the osmotic pressure difference are able to narrow the radius of the neck. Then, we consider a more realistic model by solving the complete Helfrich equation [2] including the possibility of a pulling force on the neck, e.g. due to DNA translocation through the SpoIIIE motor. For this more realistic model, the shape of the membrane neck is obtained as part of the solution.

### MINIMAL MODEL FOR THE NECK: UNIFORM CYLINDER CONNECTING TWO PLANAR MEMBRANES

We first consider a highly simplified model based on the free energy 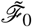 of an axisymmetric cylindrical membrane neck of radius 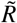 and length *L* that connects two planar membrane sheets, corresponding to the local geometry depicted in 1(b). We employ the classical Helfrich-Canham theory [2–7] for the energy of the membrane, 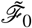, which reads

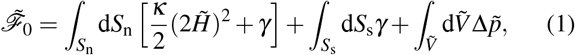

where *S_n_* and *S*_s_ are the surfaces of the membrane neck and sheets, and 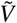 is the volume inside the neck. The tildes denote dimensional variables which will subsequently be non-dimensionalized. Here 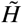 is the mean curvature, *κ* is the bending modulus, *γ* is the membrane surface tension, and 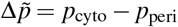 is the osmotic pressure difference. The two surrounding membrane sheets are assumed to be planar since their curvature is much less than that of the neck, thus their only contribution to the energy comes from their area times the membrane tension. In this simplified model, we neglect the bending energy of the junctions where the cylinder meets the membrane sheets. With these simplifications, the free energy 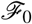 Eq. (1) reads:

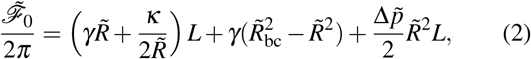

where 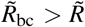 is the outer boundary condition depicted in Fig. 1(b). Minimizing Eq. (2) with respect to the radius 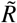 yields

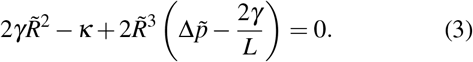

**Figure 1.**
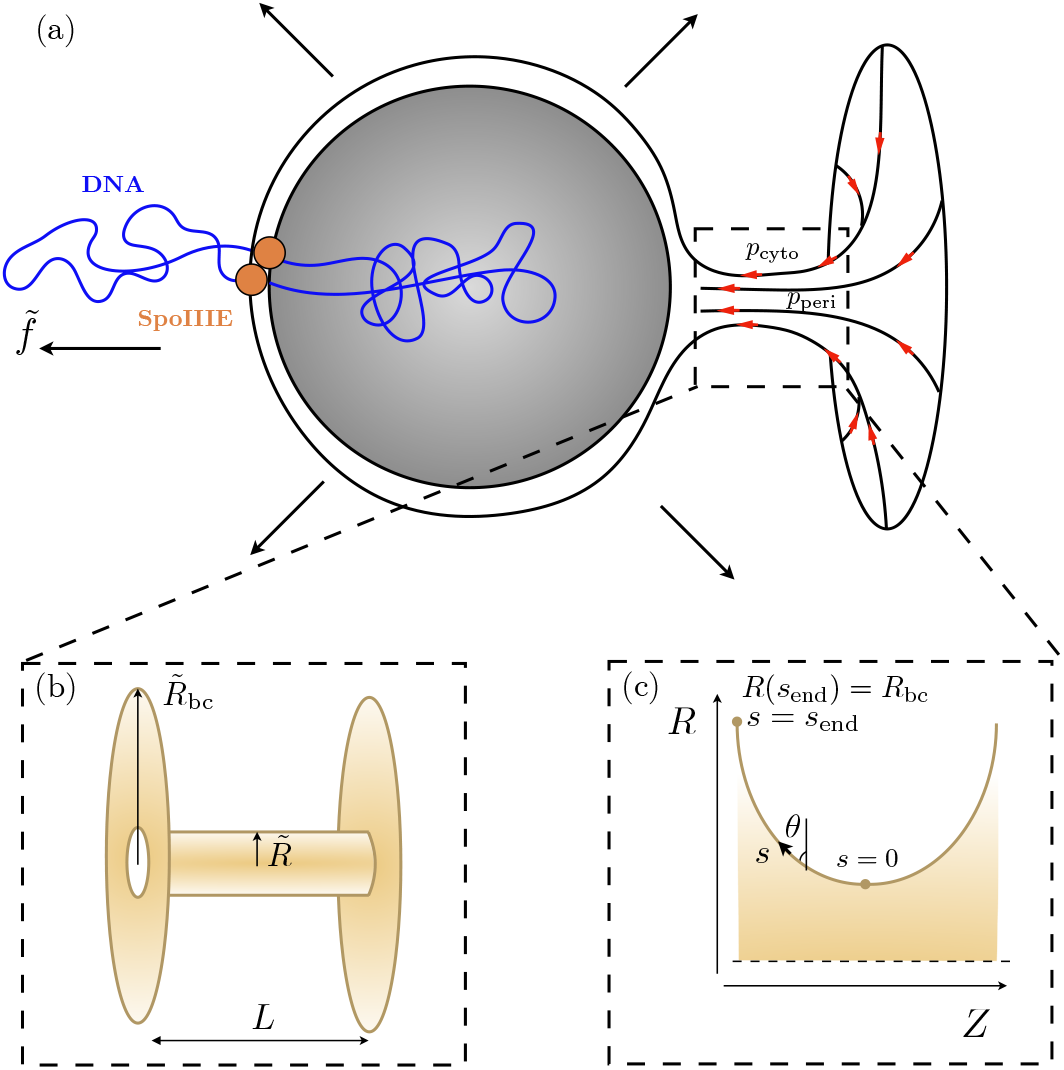
(a) Sketch of the engulfment membrane geometry during the last stage of forespore engulfment, showing the flow of lipids through the membrane neck, the osmotic pressure difference 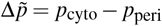 between the cytoplasm and the periplasm, and the pulling force 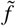 due to the SpoIIIE motors translocating DNA into the forespore. (b) Simplified model of the membrane neck, consisting of a straight cylinder of radius 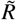 and length *L*, connecting two planar membranes of radius 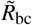. (c) Schematic of the more realistic model of the membrane neck showing the parametrization of the surface, where s is the arc length along the contour and *θ* the angle with respect to the vertical.

To non-dimensionalize Eqs. (2)-(3) we introduce the natural length scale arising from the balance between bending and tension in Eq. (1) [8] as the characteristic length scale, 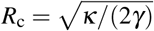, and define 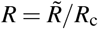. The free energy is also made dimensionless as 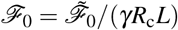. Hence, the dimensionless forms of Eqs. (2) and (3) read:

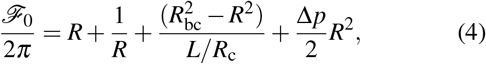

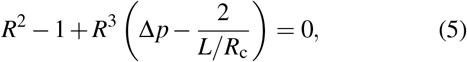

where 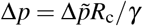.

**Figure 2.**
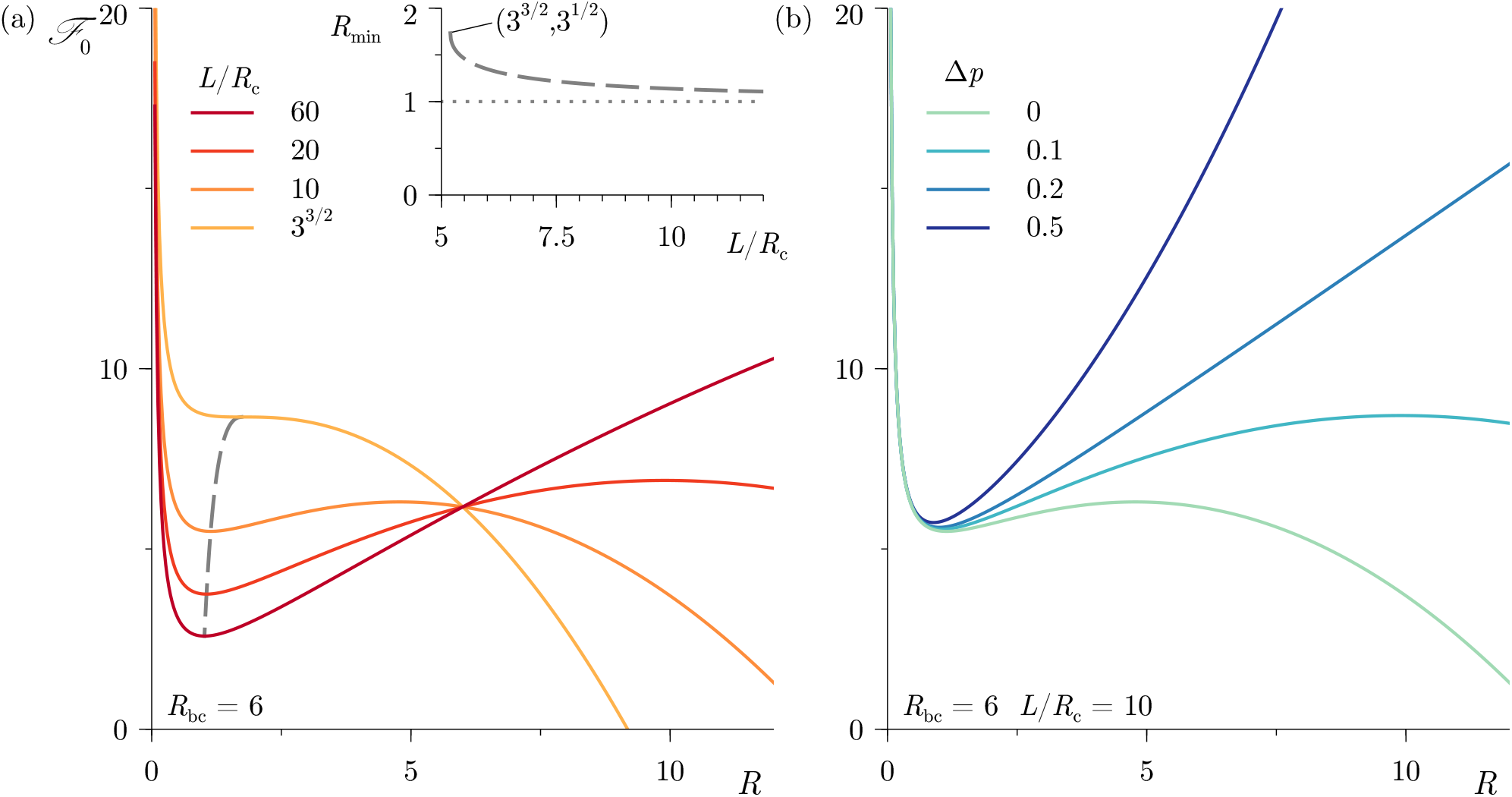
(a) Dimensionless free energy 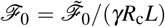 from Eq. (4) as a function of the dimensionless neck radius *R*, highlighting the existence of a local free-energy minimum provided *L*/*R*_c_ > 3^3/2^ (gray dashed curve). Here *R*_bc_ = 6 and Δ*p* = 0. The inset shows the corresponding radius *R*_min_ of the local free-energy minimum as a function of *L*/*R*_c_. (b) Same as in panel (a) but for different values of the dimensionless osmotic pressure difference Δ*p*, for a fixed length of the neck, *L*/*R*_c_ = 10.

#### Zero osmotic pressure difference

The solution of Eqs. (4) and (5) when Δ*p* = 0 reveals that for a long enough neck there is a metastable state of the neck at finite radius *R*_min_ [9, 10]. This local minimum arises from a balance between membrane bending energy, which always favors larger *R*, and membrane tension, which contributes a non-monotonic function of *R* to the membrane energy. The existence of this local minimum can be observed in Fig. 2, which shows the dimensionless free energy 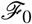 as a function of *R* for different values of *L*/*R*_c_, along with the location *R*_min_ of the local minimum. Intuitively, for *R* around *R*_min_, increasing the radius *R* increases the total amount of membrane in the vicinity of the neck and so increasing *R* is opposed by surface tension. By contrast, for larger values of *R*, increasing *R* removes more membrane from the parallel sheets (removed area scaling ~ *R*^2^) than is added to the neck (added area scaling ~ *LR*), so further increase of *R* is energetically favored (specifically when 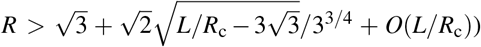. We find that, for lengths below a critical value *L/R*_c_ < 3^3/2^, the energy functional 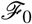 does not have a local minimum, so that expanding the radius of the neck always decreases the total energy of the system, implying the neck opens.

#### Finite osmotic pressure difference

Eq. (5) indicates that a positive osmotic pressure difference between the cytoplasm and the periplasm favors the formation of a locally stable, narrow membrane neck. Indeed, the critical neck length below which there is no locally stable narrow neck decreases as Δ*p* increases, *L/R*_c_ ≤ 2 · 3^3/2^/(2 + 3^3/2^Δ*p*). Moreover, the local maxima shown in Fig. 2(a) for Δ*p* = 0, disappears when Δ*p* > 2/(*L/R*_c_), as shown in Fig. 2(b). This implies that the free energy 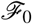 increases monotonically beyond the local minimum, i.e. the neck will not open up. Moreover, the minimum radius of the neck *R*_min_ always decreases as Δ*p* increases.

### EQUILIBRIUM SHAPES OF THE AXISYMMETRIC MEMBRANE NECK

We now go beyond the minimal model described in the previous section by solving the complete Helfrich equation, where the shape of the neck is obtained as part of the solution. To this end, we describe the equilibrium shape of an axisymmetric and mirror symmetric membrane neck connecting two membrane sheets, corresponding to the local geometry of the neck connecting the forespore engulfing and mother cell membranes (see Figs. 1a and c). The energy functional 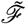 consists of a term accounting for membrane bending and tension, 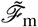, another term accounting for the osmotic pressure pressure difference between cytoplasm and periplasm, 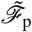, and a term representing any pulling force *f* on the forespore, 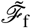[2–8, 11, 12]:

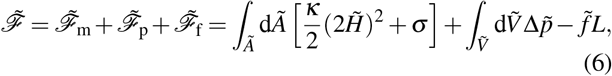

where 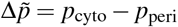, 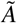 is the area of the membrane neck, and 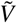 is the volume inside the neck.

**Figure 3.**
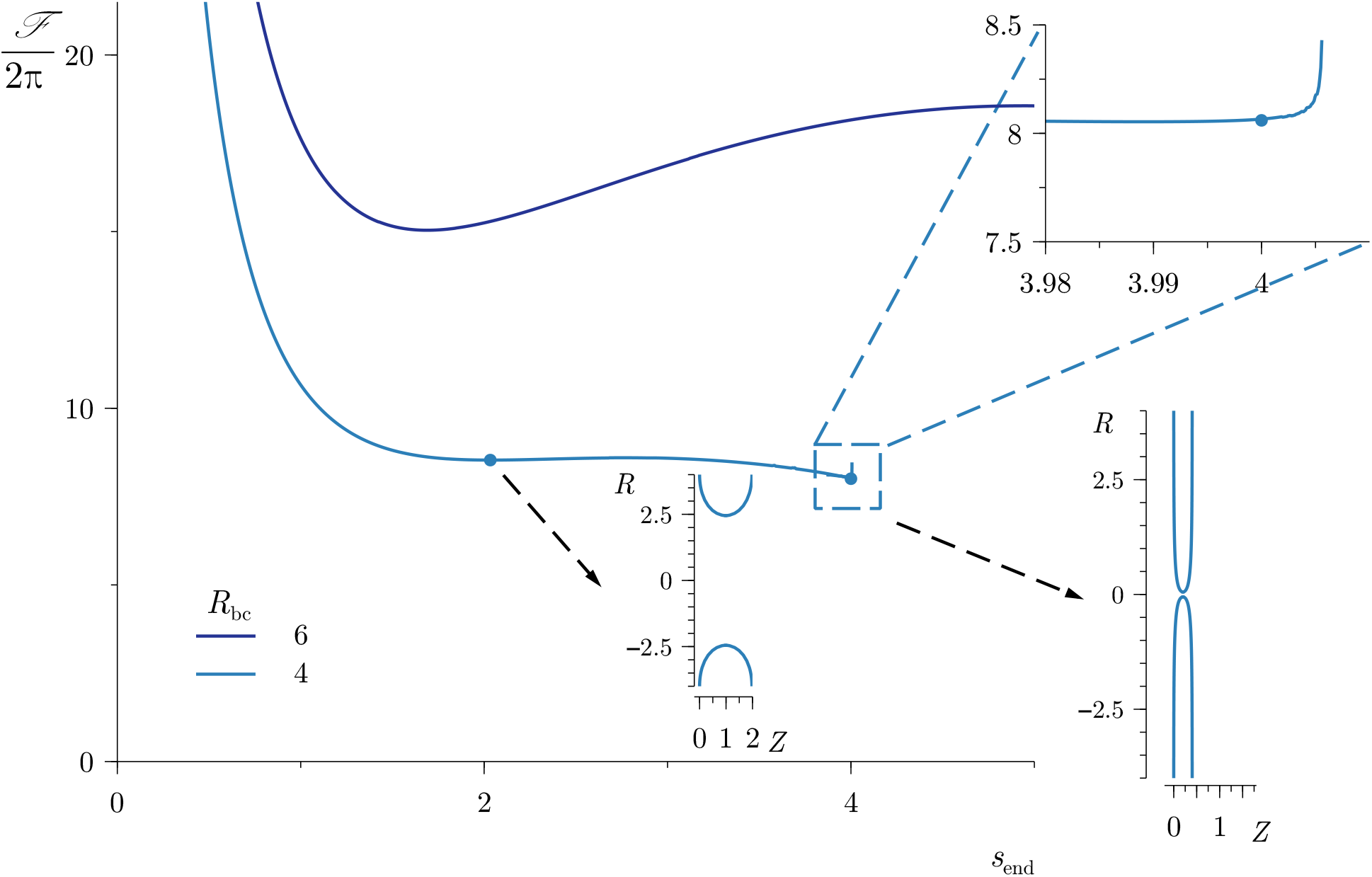
Dimensionless free energy 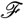 from Eq. (6) as a function of the arc length *s*_end_ for different values of the radial boundary size *R*_bc_. The insets show shapes of the neck for *R*_bc_ = 4 and two different values of *s*_end_, corresponding to two local minima. The zoomed-in region depicts the sudden increase of free energy as the neck narrows.

For axisymmetric surfaces, it proves convenient to describe the surface in terms of the angle *θ*(*s*) between the contour and the ordinate axis, as parametrized by the arclength s (see sketch in Fig. 1c). To non-dimensionalize Eq. (6), we again introduce the natural length scale 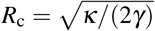 as the characteristic length. The definition of the radial *R*(*s*) and axial *Z*(*s*) coordinates, and the first variation of Eq. (6) yield the following nonlinear boundary-value problem [11, 13, 14]

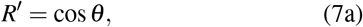

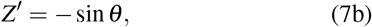

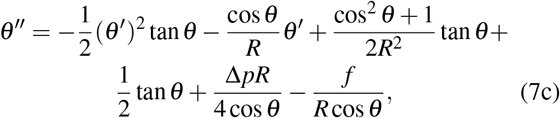

where derivatives are with respect to contour length *s*, 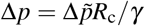, and 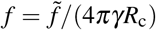. Concerning the boundary conditions, we impose *R*(*s* = *s*_end_) = *R*_bc_, *Z*(*s* = *s*_end_) = 0, *θ*(*s* = *s*_end_) = 0, and *θ*(*s* = 0) = *π*/2. The dimensionless free energy of the system is computed as

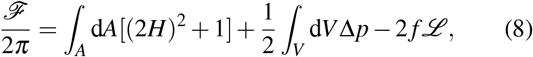

where 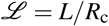. The system of differential equations (7a)-(7c) is solved using the Matlab® built-in boundary-value problem solver bvp4c [15, 16].

Fig. 3 shows the dimensionless version of the free energy from Eq. (6), as a function of the arc length *s*_end_ for different values of the radial boundary size Rbc indicated in the legend. When *R*_bc_ ≳ 3, the free energy exhibits two local minima. One of them corresponds to a relatively small value of *s*_end_, where the minimum neck radius is large, i.e. the neck opens up. The other local minimum occurs at a larger value of *s*_end_ with a larger associated free energy, corresponding to a narrow neck.

Increasing the value of send beyond the latter local minimum increases the total free energy drastically due to a sharp increase of the bending energy, as shown for *R*_bc_ ≲ 4 (zoomed region in Fig. 3). When R_bc_ < 3, the free energy only displays one local minimum, which corresponds to that of a narrow neck. Fig. 4 is obtained using the same values of the parameters as in Fig. 3, but displaying some of the multiple higher-energy branches arising for sufficiently large values of *s*_end_.

**Figure 4.**
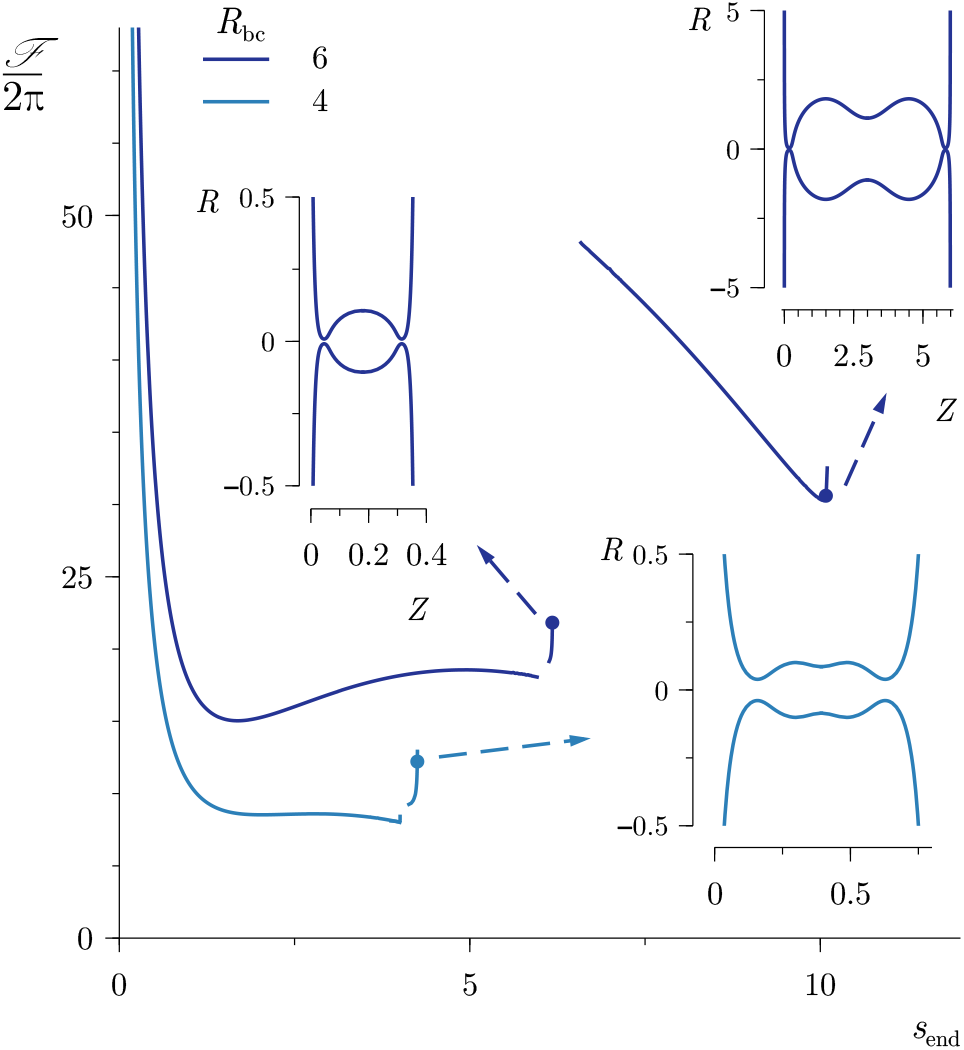
Same as in Fig. 3 but displaying some of the multiple branches arising for sufficiently large values of *s*_end_.

Additionally, Fig. 5 shows the role of the pulling force *f* and the osmotic pressure difference Δ*p* on the free energy and the shape of the membrane neck. Panel (a) depicts the dimensionless free energy 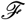 as a function of *s*_end_ for *f* = 0 and *f* = 1.35 (see Table I), and Δ*p* = 0. The insets, showing the shape of the axisymmetric membrane neck at different values of send, demonstrate that, for the same values of Rbc and send, the pulling force tends to increase both the length of the neck and the minimum neck radius. Panel (b) displays the free energy with a non-zero value of the pulling force and the osmotic pressure difference, i.e. *f* = 1.35 and Δ*p* = 0.5. As expected, incorporating a non-zero value of the osmotic pressure difference between the cytoplasm and the periplasm, slightly favors the narrowing of the membrane neck.

**Table I.** Estimates of the physical and dimensionless parameters

| Parameter | Value |
| --- | --- |
| $\kappa$ | $20 \text{ k}_B T$ [8] |
| $\gamma$ | $0.1\text{--}10 \text{ pN nm}^{-1}$ [17–19] |
| $L$ | $40 \text{ nm}$ |
| $\Delta\tilde{p}$ | $60 \text{ kPa}$ [20] |
| $\tilde{f}$ | $50\text{--}100 \text{ pN}$ [21, 22] |
| $R_c = \sqrt{\kappa/(2\gamma)}$ | $2\text{--}20 \text{ nm}$ |
| $L/R_c$ | $9\text{--}90$ |
| $\Delta p = \Delta\tilde{p}R_c/\gamma$ | $0.012\text{--}12$ |
| $f = \tilde{f}/(4\pi\gamma R_c)$ | $0.4\text{--}4$ |

We now estimate some of the physical and dimensionless parameters used in the model and show that realistic values of the membrane tension and the osmotic pressure difference are able to sufficiently narrow the neck radius, to allow FisB proteins to interact and accumulate inside the neck. These estimates are shown in Table I. In particular, we consider a pulling force of 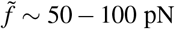, which corresponds to the SpoIIIE maximum stall force [21, 22], and an osmotic pressure difference of 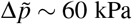, as estimated in Ref. [20]. Based on previous works [1, 23], we assume that FisB proteins are able to undergo homo-oligomerization in trans when the radius of the membrane neck is below ~ 10 nm, which is within the range of the estimated characteristic neck radius shown in Table I, 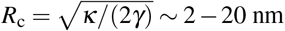 nm, where we have considered *κ* ~ 20 *k_B_T* [8], and *γ* ~ 0.1 — 10 pN nm^−1^ [17–19] (see last section of this Appendix for an upper estimate of the membrane tension). According to the model, when Δ*p* = *f* = 0, for the local minimum of the free energy 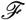 corresponding to the larger value of *s*_end_, shown in Fig. 3, the dimensional minimum radius of the neck is 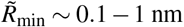, which is significantly below the minimum radius required for FisB to interact. Moreover, the local free-energy minimum corresponding to a wider neck yields a minimum neck radius 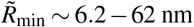, implying that, if membrane tension increases sufficiently, the neck radius for this local minimum can also be narrow enough for FisB to interact and accumulate inside the neck.

When *f* = 1.35 and Δ*p* = 0, which corresponds to, for instance, *γ* = 0.84pN nm^−1^ and 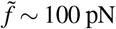, we find 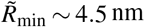, corresponding to the membrane neck shown in the right bottom inset of Fig. 5(a). Hence, the action of a pulling force increases the radius of the neck, as compared to the case of *f* = 0 shown in Fig. 3 and 5(a). If we now consider the case of a non-zero value of the osmotic pressure difference, shown in Fig. 5(b), where we have used *f* = 1.35 and Δ*p* = 0.5, corresponding to *γ* ~ 0.84 pN nm^−1^, 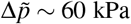, and 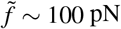, the minimum radius is 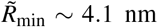, thereby favoring the accumulation of FisB in the neck. Nonetheless, such osmotic pressure difference only reduces the minimum neck radius slightly, as we also observed in our highly simplified model shown in Fig. 2(b).

### FLUX OF LIPIDS

To estimate the membrane tension difference between the two ends of the neck that would drive a flux of lipids from the mother cell to the forespore of ~ 10^3^ lipids *s*^−1^ (see *Forespore inflation accompanies membrane fission* in the main text), we balance the force due to the membrane tension difference Δ*γ* and the drag force on a cylinder (i.e. the neck in our highly simplified model), translating parallel to its axis in an outer bath of viscosity *μ*,

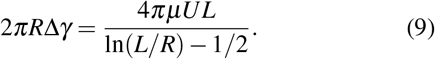

Considering that 1 lipid occupies ~ 0.7 nm^2^ and assuming a neck radius of *R* ~ 5 nm, the required lipid velocity along the neck is *U* ~ 22.3 nm *s*^−1^. Taking *L* ~ 40 nm and *μ* ~ 10^-3^ Pa s, the estimated membrane tension difference is Δ*γ* ~ 2.3 × 10^-7^ pN nm^-1^. Hence, a flux of ~ 10^3^ lipids s^-1^ would require a relatively small excess membrane tension in the forespore engulfing membrane compared to the mother cell membrane, which we assume is of the same order as the estimate in Table I [17–19]. So in the absence of an accumulation of FisB in the neck that would slow lipid flow, a flux of lipids in the neck could readily provide ~ 10^3^ lipids s^-1^ to the forespore membranes.

**Figure 5.**
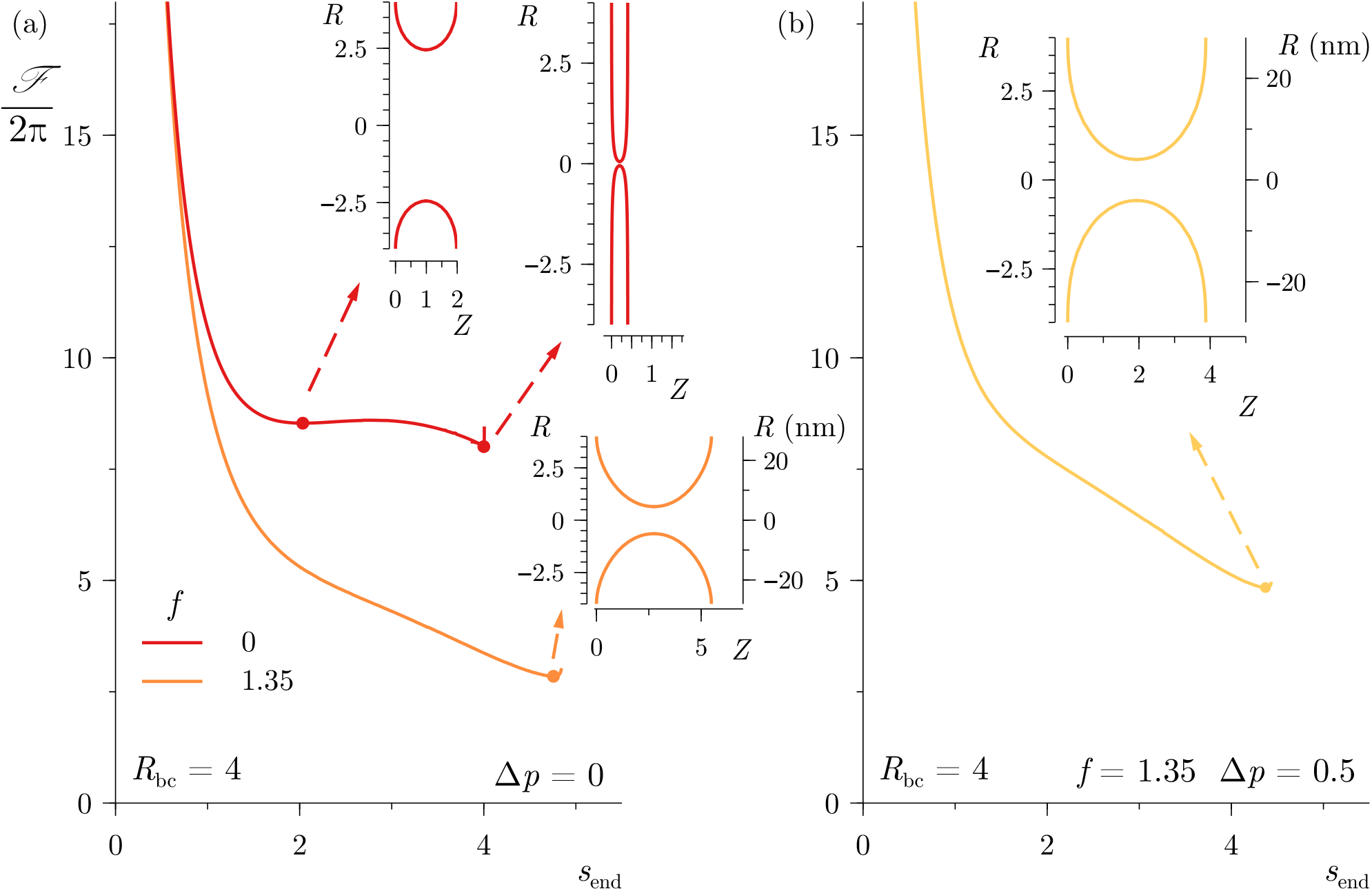
Dimensionless free energy 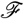 from Eq. (6) as a function of the arc length *s*_end_ for *R*_bc_ = 4. Panel (a) shows 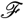 for two different values of the dimensionless pulling force *f*, with Δ*p* = 0, and panel (b) shows the case *f* = 1.35, Δ*p* = 0.5. The insets show the shapes of the neck for different values of *s*_end_ indicated with dots.

### COMPARING THE STALL FORCE OF SPOIIIE TO THE OSMOTIC PRESSURE DIFFERENCE

Here we analyze how the osmotic pressure difference between the mother cell and the forespore compares to the stall/disassembly force of SpoIIIE [21, 22]. According to Ref. [20], the excess pressure in the forespore is of the order of 60 kPa, which can be related to a salt concentration difference Δ*ρ* between the mother cell and the forespore via Δ*p* = *N_A_k_B_T*Δ*ρ*, yielding Δ*ρ* ~ 25 mM. Taking the salt concentration in the mother cell to be *ρ*_mc_ ~ 200 mM [24, 25], the salt concentration inside the forespore is *ρ*_fs_ ~ 225 mM. We can now estimate the energy required to move one salt ion into the forespore as *E* = *k_B_T*log(*ρ*_fs_/*ρ*_mc_) ~ 2.23kBT. This estimate is of the same order as the energy available from SpoIIIE, which is its stall maximal force, ~ 50 pN [21, 22], multiplied by half the base spacing, i.e. ~ 0.17 nm, since each translocated base of DNA brings ~2 counterions, which yields *E*_SpoIIIE_ ~ 2.1*k*_B_*T*. This near equality of energies suggests that SpoIIIE works near its stall force, by creating the highest pressure difference it can in the forespore.

Moreover, we can estimate the forespore membrane tension that would be required to balance this osmotic pressure difference (for this estimate, we neglect the possible role of the peptidoglycan surrounding the forespore in opposing this osmotic pressure difference). Assuming that the forespore is a sphere surrounded by two membranes, the equilibrium excess pressure is given by the Laplace law, Δ*p* = 4*γ*/*R*_fs_. Taking *R*_fs_ ~ 400 nm, we obtain *γ* ~ 6 pN nm^-1^. According to Refs. [17, 19], the rupture tension of a lipid membrane is γ_rup_ ~ 20 pN nm^-1^. Nonetheless, as shown in Ref. [18], the membrane rupture tension depends on the application time of the tension, and it can be of the order of *γ*_rup_ ~ 2 – 30 pN nm^-1^ when *γ*_app_ ~ 4 – 0.01 min. Altogether, we argue that forespore inflation can give rise to an increase of membrane tension enough to narrow the neck below 10 nm (see Equilibrium shapes of the axisymmetric membrane neck). Moreover, given that the characteristic time scale of forespore inflation is *t* ~ 1 – 10 min, the increased membrane tension due to forespore inflation may contribute directly to membrane fission.

